# Precise transcription timing by a second-messenger drives a bacterial G1/S cell cycle transition

**DOI:** 10.1101/675330

**Authors:** Andreas Kaczmarczyk, Antje M. Hempel, Christoph von Arx, Raphael Böhm, Badri N. Dubey, Jutta Nesper, Tilman Schirmer, Sebastian Hiller, Urs Jenal

## Abstract

Bacteria adapt their growth rate to their metabolic status and environmental conditions by modulating the length of their quiescent G1 period. But the molecular mechanisms controlling G1 length and exit from G1 are poorly understood. Here we identify a key role for the second messenger c-di-GMP, and demonstrate that a gradual increase in c-di-GMP concentration determines precise gene expression during G1/S in *Caulobacter crescentus*. We show that c-di-GMP strongly stimulates the kinase ShkA, activates the TacA transcription factor, and initiates a G1/S-specific transcription program leading to cell morphogenesis and S-phase entry. C-di-GMP activates ShkA by binding to its central pseudo-receiver domain uncovering this wide-spread domain as a novel signal input module of bacterial kinases. Activation of the ShkA-dependent genetic program also causes c-di-GMP to reach peak levels, which triggers S-phase entry and, in parallel, promotes proteolysis of ShkA and TacA. Thus, a gradual increase of c-di-GMP results in a precisely tuned ShkA-TacA activity window enabling G1/S specific gene expression before cells commit to replication initiation. By defining a regulatory mechanism for G1/S control, this study contributes to understanding bacterial growth control at the molecular level.

**GRAPHICAL ABSTRACT:** 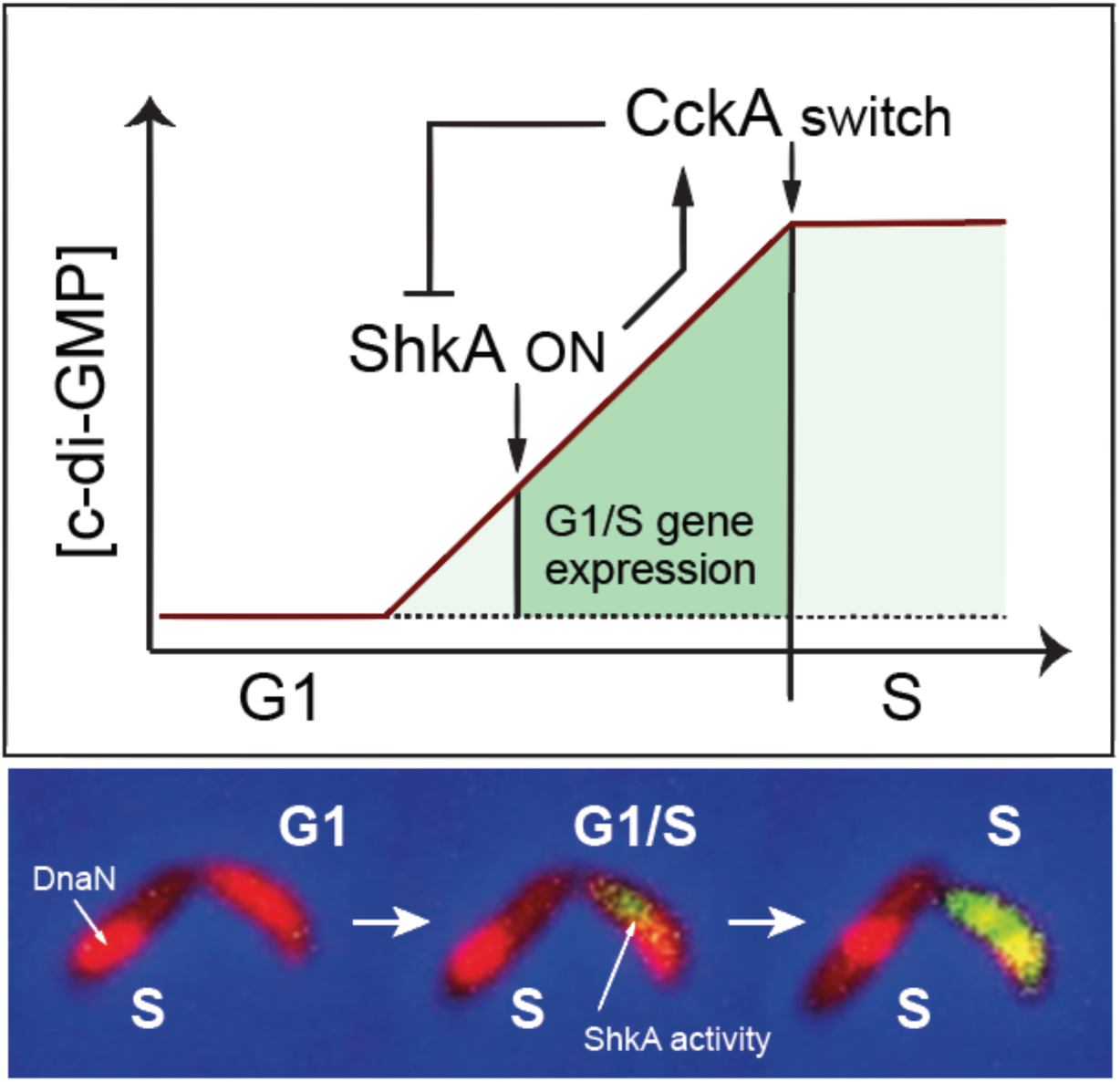

## INTRODUCTION

The bacterial cell cycle is divided into three periods: after cell division and before initiation of chromosome replication (B or G1); chromosome replication (C or S); and cell division (D or G2) ^1^. Since chromosome replication and cell division (C and D periods) remain constant over a wide range of growth rates ^2,3^, the step committing cells to initiate chromosome replication largely determines bacterial proliferation rates. Bacteria like *Escherichia coli* or *Bacillus subtilis* can increase their growth rate by bypassing the B period and by initiating replication multiple times per division cycle ^2^. In contrast, *Caulobacter crescentus* strictly separates its cell cycle stages. An asymmetric division generates a sessile stalked (ST) cell, which directly re-enters S-phase, and a motile swarmer (SW) cell that remains in G1 for a variable time depending on nutrient availability ^4,5^. Coincident with G1/S transition, motile SW cells undergo morphogenesis to gain sessility (Fig. 1a). But what determines the length of G1 in this organism has remained unclear.

**Fig. 1:**
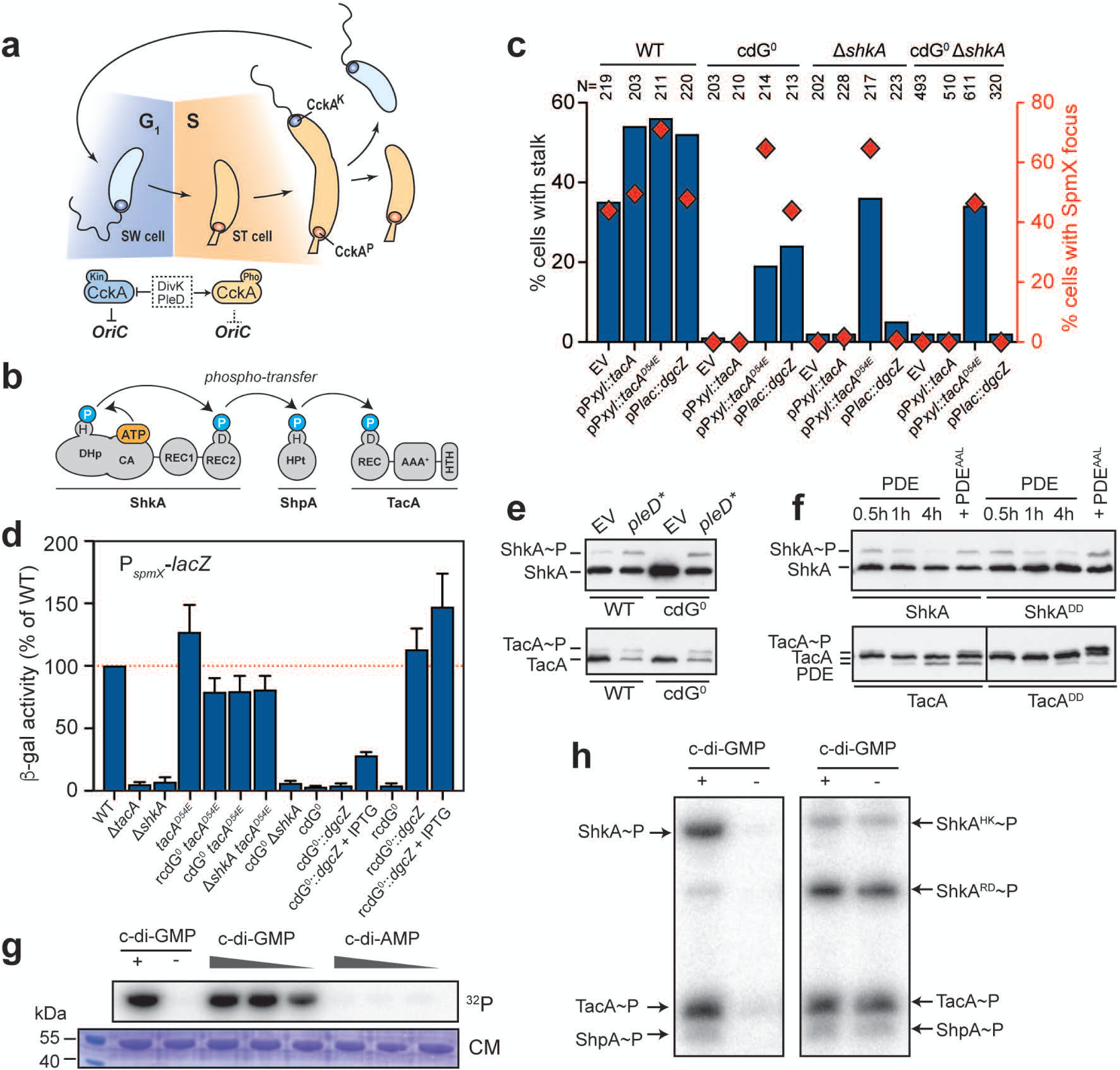
C-di-GMP controls the ShkA-TacA phosphorelay *in C. crescentus*. **a**, Schematic of the *C. crescentus* cell cycle with swarmer (SW) and stalked (ST) cells colored in blue and orange, respectively, and the G1- and S-phases of the cell cycle indicated in similar colors. Stage-specific kinase (Kin) and phosphatase (Pho) activities of the cell cycle kinase CckA are indicated below with the coloring referring to stage-related activities. The response regulators DivK and PleD that control the CckA switch are highlighted. **b**, Schematic of the ShkA-ShpA-TacA phosphorelay. **c**, Quantification of cells with stalks and SpmX-mCherry foci of strains expressing a chromosomal *spmX-mCherry* fusion and plasmid-driven *tacA*, *tacA^D54E^* or the heterologous diguanylate cyclase *dgcZ*. EV, empty vector control. The number of cells analyzed (N) is indicated above the graph. For representative micrographs, see Supplementary Fig. 1. **d**, Activity of the *spmX* promoter in the indicated strains harboring a P*spmX*-*lacZ* transcriptional reporter (plasmid pRKlac290-*spmX*). IPTG indicates induction of *dgcZ* expression from P*lac* by addition of 300 µM IPTG. Shown are mean values and standard deviations (N>3). **e**, Phos-tag PAGE immunoblots of strains producing 3xFLAG-tagged ShkA or TacA from the native chromosomal loci and a plasmid expressing the constitutively active diguanylate cyclase PleD*. **f**, Phos-tag PAGE immunoblots of strains encoding 3xFLAG-tagged ShkA or TacA at the native loci or the respective mutant alleles encoding ClpXP protease resistant variants TacA^DD^ or ShkA^DD^. To manipulate c-di-GMP levels, all strains carried a plasmid expressing wild-type or a catalytic mutant (AAL) of the phosphodiesterase (PDE) PA5295 from a cumate-inducible promoter. Samples were harvested at the time indicated after addition of 100 µM cumate to the medium. Control samples (PA5295^AAL^) were harvested 4 hours after induction. **g**, *In vitro* phosphorylation assays with purified ShkA in the presence of different concentrations of nucleotides (1 mM, 100 µM, 10 µM). Positive and negative controls contained 100 µM c-di-GMP and ddH_2_O, respectively. Reactions were initiated by addition of 500 µM radiolabeled ATP and allowed to proceed for 15 minutes at room temperature. **h**, *In vitro* phosphorylation assays with purified components (4 µM) of the ShkA-ShpA-TacA phosphorelay with (+) or without (-) c-di-GMP (76 µM). ShkA^HK^ or ShkA^RD^ denote truncated ShkA constructs that either constitute the kinase catalytic core plus REC1 (ShkA^HK^) or REC2 (ShkA^RD^). Reactions were initiated by addition of 500 µM ATP and allowed to proceed for 15 minutes at room temperature.

In *C. crescentus*, replication initiation is regulated by the cell cycle kinase CckA. CckA is a bifunctional enzyme that acts as a kinase for the replication initiation inhibitor CtrA in G1, but switches to being a phosphatase in S phase, resulting in the inactivation of CtrA and clearance of the replication block ^6^ (Fig. 1a). The CckA switch is governed by two response regulators, DivK and PleD. While DivK controls CckA activity through protein-protein interactions ^7–9^, PleD is a diguanylate cyclase, which is responsible for the characteristic oscillation of c-di-GMP during the cell cycle ^10^. The concentration of c-di-GMP, below the detection limit in G1, increases during G1/S to reach peak levels at the onset of S-phase ^11^ where it allosterically stimulates CckA phosphatase ^9^ (Fig. 1a).

Activation of DivK and PleD is directed by the SpmX scaffolding protein, which accumulates during G1/S and recruits DivJ, the kinase of DivK and PleD, to the incipient stalked cell pole ^12^. This makes SpmX accumulation the earliest known event to trigger S phase entry. However, it is unclear how *spmX* expression is timed during G1/S to initiate replication. Transcription of *spmX* is regulated by the response regulator TacA, which in its phosphorylated form also induces the expression of a large set of genes required for SW-to-ST cell morphogenesis ^12,13^. TacA is activated via a multi-step phosphorylation cascade (Fig. 1b) consisting of the sensor histidine kinase ShkA, and the phosphotransferase protein ShpA ^12,13^. ShkA is a multidomain protein kinase with a catalytic domain (CA) that binds ATP and transfers a phosphate via the conserved His residue of the dimerization histidine phosphotransfer domain (DHp) to a conserved Asp residue of the C-terminal receiver domain (REC2) ^12,13^ (Fig. 1b). Thus, this phosphorelay system controls both replication initiation and stalk biogenesis. However, the signal activating ShkA has remained unknown (Fig. 1b).

Here we demonstrate that an upshift in levels of the second messenger c-di-GMP determines precise gene expression during G1/S. A combination of genetic, biochemical and cell biology data revealed that c-di-GMP controls the ShkA-TacA pathway by directly binding to the ShkA sensor histidine kinase and strongly stimulating its kinase activity. We demonstrate that binding of c-di-GMP to a pseudo-receiver domain of ShkA abrogates ShkA auto-inhibition leading to the activation of the ShkA-TacA pathway. C-di-GMP-mediated activation of ShkA and the subsequent c-di-GMP-mediated proteolysis of ShkA together define a window of ShkA activity during the cell cycle. This window is sharpened to a narrow, G1/S-specific period by TacA degradation, which precedes the degradation of ShkA. Thus, the exact timing of G1/S-specific gene expression results from the consecutive c-di-GMP-mediated activation and degradation of ShkA-TacA phosphorylation components.

## RESULTS

### C-di-GMP stimulates ShkA kinase activity to control the ShkA-TacA phosphorylation cascade

A *C. crescentus* strain lacking all diguanylate cyclases (cdG^0^ strain), which was generated to remove all traces of the second messenger c-di-GMP, shows strong developmental and morphological abnormalities with mutant cells being irregularly shaped, elongated and lacking all polar appendages ^11^. Because stalk biogenesis depends on an active ShkA-TacA phosphorelay (Fig. 1b) ^13^, we tested if c-di-GMP controls the ShkA-TacA pathway. Expression of TacA^D54E^, a phospho-mimetic form of TacA ^13^, restored stalk biogenesis, transcription of the TacA targets *staR* and *spmX*, as well as localization of a SpmX-mCherry fusion to the stalked pole (Fig. 1c,d; Supplementary Fig. 1). *spmX* and *staR* transcription was also restored when c-di-GMP was reintroduced by expression of the heterologous diguanylate cyclase (DGC) *dgcZ* from *E. coli* in the cdG^0^ background or in a strain that lacks all DGCs and phosphodiesterases (PDE) (rcdG^0^ strain) (Fig. 1d; Supplementary Fig. 1). Phos-tag PAGE analysis revealed that TacA and ShkA were unphosphorylated in the cdG^0^ strain. This phosphorylation was restored upon expression of the constitutively active DGC PleD* ^14^ (Fig. 1e). Likewise, expression of the heterologous phosphodiesterase PA5295 from *Pseudomonas aeruginosa*, which also lowers c-di-GMP levels, in wild-type cells reduced TacA and ShkA phosphorylation (Fig. 1f). These results indicate that c-di-GMP acts upstream of and is required for the activity of the ShkA-TacA phosphorelay.

*In vitro* phosphorylation assays with purified ShkA or with all components of the phosphorelay demonstrated that c-di-GMP strongly and specifically stimulates ShkA autokinase activity (Fig. 1g,h; Supplementary Fig. 2a). Moreover, purified ShkA bound c-di-GMP with a KD in the sub-micromolar range (Supplementary Fig. 2b). Altogether, we conclude that c-di-GMP activates the ShkA-TacA pathway by directly binding to and allosterically stimulating ShkA kinase.

### C-di-GMP activates ShkA by binding to the REC1 pseudo-receiver domain

We next sought to dissect the mechanism of c-di-GMP-mediated ShkA activation. We devised a genetic selection (see Materials and Methods) to isolate *shkA* mutations that restored *spmX* expression in a rcdG^0^ background. Independent mutations were identified in two residues (D369, R371) within a short stretch of three highly conserved amino acids in the linker region between REC1 and REC2 (hereafter referred to as the “DDR” motif) (Fig. 2a; Supplementary Fig. 3). These results identified the REC1-REC2 linker region as a critical determinant of ShkA activation.

**Fig. 2:**
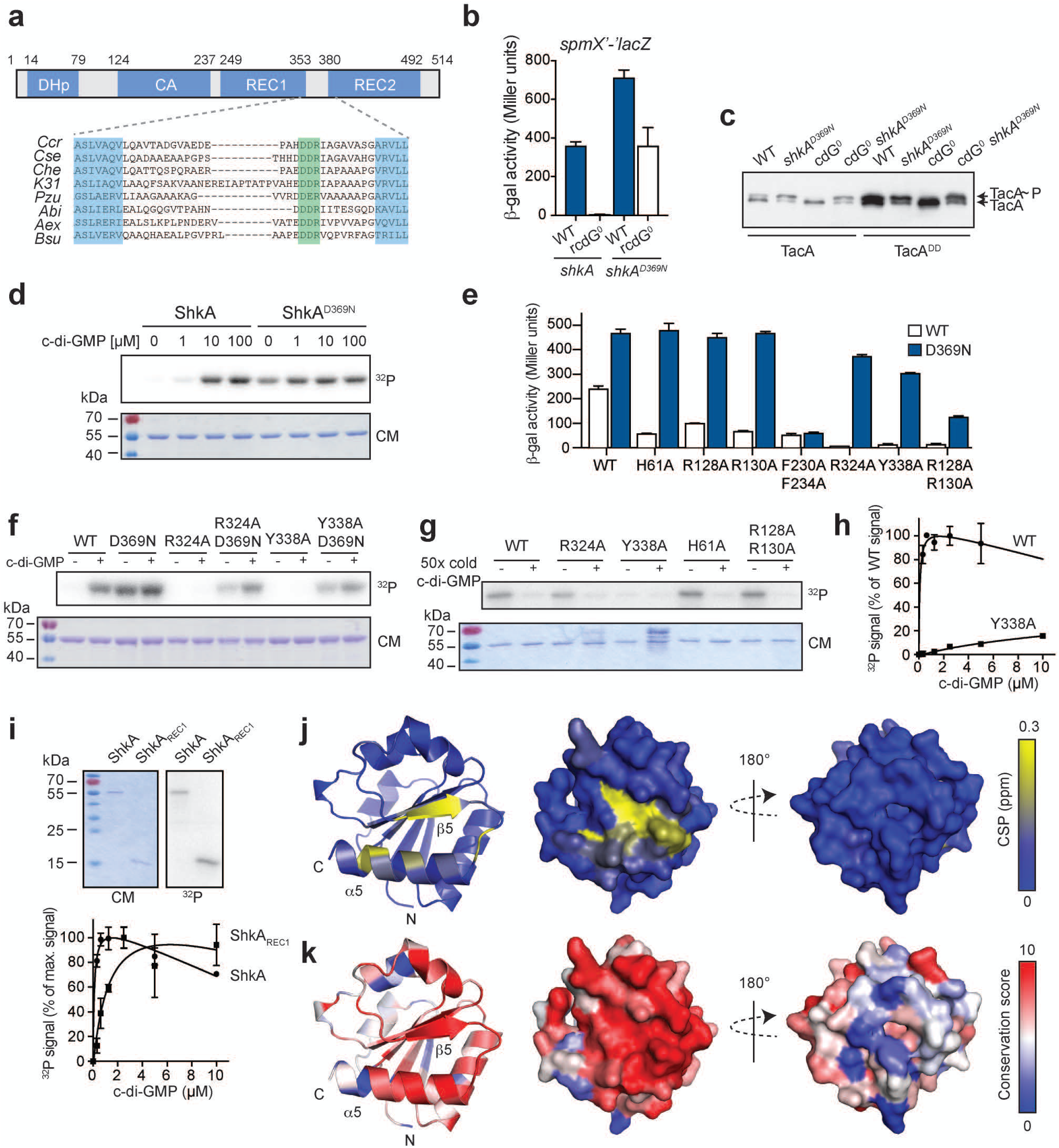
C-di-GMP activates ShkA by binding to the REC1 pseudo-receiver domain. **a**, Schematic of ShkA domain architecture approximately drawn to scale (top) and alignment of the REC1-REC2 linker harboring the DDR motif (highlighted in green) of ShkA orthologs from several *Alphaproteobacteria* (bottom). Ccr, *C. crescentus*; Cse, *Caulobacter segnis*; Che, *Caulobacter henricii*; K31, *Caulobacter sp*. K31; Pzu, *Phenylobacterium zucineum*; Abi; *Asticcacaulis biprosthecium*; Aex, *Asticcaucaulis excentricus*; Bsu, *Brevundimonas subvibrioides*. The dimerization histidine phosphotransfer (DHp), the Catalytic ATP binding (CA) and the receiver domains (REC1, REC2) are indicated. **b**, Activity of the *spmX* promoter in indicated strains harboring the *spmX’-‘lacZ* reporter fusion (plasmid pAK502-*spmX*). The *shkA^D369N^* allele is expressed from the native chromosomal locus. Mean values and standard deviations are shown (N=3). **c**, Phos-tag PAGE immunoblots of indicated strains producing 3xFLAG-tagged TacA or TacA^DD^. The *tacA^DD^* allele is expressed from the native chromosomal locus and blots are probed with anti-FLAG antibodies. **d**, *In vitro* autophosphorylation assays of ShkA and ShkA^D369N^. Reactions contained 5 µM ShkA and different concentrations of c-di-GMP as indicated. Upon addition of radiolabeled ATP, autophosphorylation was allowed to proceed for 3.5 minutes at room temperature. Top: autoradiograph; bottom: Coomassie stain of the same gel. **e**, Activity of the *spmX* promoter in the Δ*shkA* mutant harboring a *spmX’-‘lacZ* reporter fusion (plasmid pAK502-spmX) and expressing different *shkA* alleles in trans from plasmid pQF with the indicated amino acid substitutions alone (WT, white bars) or in combination with the D369N substitution (blue bars). Shown are mean values and standard deviations (N=3). **f**, *In vitro* autophosphorylation assays of wild-type ShkA and indicated mutant variants with (10 µM) or without c-di-GMP. Autophosphorylation was allowed to proceed for 5 min at room temperature. Top: autoradiograph; bottom: Coomassie stain of the same gel. **g**, Autoradiographs of purified ShkA and ShkA mutant variants (0.5 µM) UV-crosslinked with 10 µM [^32^P]c-di-GMP with or without addition of a 50-fold molar excess of non-labeled c-di-GMP. Top: autoradiograph; bottom: Coomassie stain of the same gel. **h**, Quantified autoradiographs of purified ShkA and the ShkA^Y338A^ variant (0.5 µM) UV-crosslinked with increasing concentrations of [32P]c-di-GMP. Shown are mean values and standard deviations (N=2). **i**, Autoradiographs and Coomassie stain of the same gel of purified ShkA and the isolated REC1 domain (ShkAREC1) (0.5 µM) after UV-crosslinking with 10 µM [^32^P]c-di-GMP (top). Quantified autoradiographs of purified ShkA and ShkAREC1 (0.5 µM) after UV-crosslinking with increasing concentrations of [^32^P]c-di-GMP (bottom). Mean values and standard deviations are shown (N=2). **j**, Cartoon and surface representation of the ShkAREC1 homology model with NMR chemical shift perturbations (CSPs) upon c-di-GMP binding indicated by a blue-to-yellow gradient. **k**, Conservation score of ShkA orthologs (see Materials and Methods).

We further characterized the D369N variant, and confirmed that TacA phosphorylation levels and *spmX* transcription were restored to wild-type levels in the rcdG^0^ background harboring the *shkA^D369N^* allele (Fig. 2b,c). Also, purified ShkA^D369N^ protein showed strong autophosphorylation activity even in the absence of c-di-GMP (Fig. 2d). The mutant retained its ability to bind c-di-GMP *in vitro* (Supplementary Fig. 2c) and could still be partially stimulated by c-di-GMP both *in vivo* (Fig. 2b) and *in vitro* (Supplementary Fig. 2d). Thus, mutations in the DDR motif uncouple ShkA activity from c-di-GMP without interfering with c-di-GMP binding.

We used the D369N ShkA variant to identify residues involved in c-di-GMP-dependent activation. Mutations specifically interfering with c-di-GMP binding or with c-di-GMP-dependent activation should be rescued when combined with D369N, while more general kinase defects would not be recuperated. An alignment of ShkA orthologs from *C. crescentus* and related organisms revealed a total of 25 candidate residues for c-di-GMP binding distributed throughout the entire protein (Supplementary Fig. 3). Of all the mutants that severely affected ShkA activity (Supplementary Fig. 4a,b) and that were rescued *in vivo* (Fig. 2e) and *in vitro* (Fig. 2f) when combined with D369N, only one, Y338A, in REC1, interfered with c-di-GMP binding (Fig. 2g,h).

Thus, Y338 in REC1 is likely part of the c-di-GMP binding site. Indeed, the purified REC1 domain alone was able to bind c-di-GMP, although with lower affinity than full-length ShkA (Fig. 2i). NMR spectroscopy with REC1 revealed a fold reminiscent of prototypical REC domains, except that helix α3 is not present irrespective of the presence or absence of c-di-GMP (Supplementary Fig. 4c-f). Upon addition of c-di-GMP, REC1 shows chemical shift perturbations (CSPs), which cluster on the α4-β5-α5 surface (Fig. 2j; Supplementary Fig. 4g). Importantly, these residues include Y338 and are well conserved (Fig. 2k; Supplementary Fig. 3). Some of the residues implicated in c-di-GMP binding by NMR were also important for ShkA activity *in vivo* (Supplementary Fig. 4h). Altogether, these experiments revealed a REC1 pseudo-receiver domain of ShkA as the primary docking site for c-di-GMP, and identified residues directly involved in c-di-GMP binding and c-di-GMP-mediated activation of ShkA. *In silico* analysis revealed that pseudo-receiver domains, although often not annotated, are widespread among histidine kinases (Supplementary Fig. 5, Supplementary Data 1). We propose that pseudo-receiver domains have lost their original phosphotransfer function but during evolution have adopted novel signaling functions and may represent a large class of novel kinase input domains.

Similar to the two identified DDR mutants, the kinase activity of ShkA is also uncoupled from c-di-GMP when conserved residues of REC2 are mutated (Supplementary Fig. 2e) or when the C-terminal REC2 domain is removed from the catalytic core (DHp, CA and REC1 domain) (Fig. 1h). Thus, the REC2 domain and the DDR linker motif inhibit ShkA autophosphorylation, maintaining it in an inactive state. We propose that c-di-GMP binding to REC1 overrides this auto-inhibition and activates the enzyme.

### C-di-GMP defines a narrow window of ShkA-TacA activity during G1/S

The c-di-GMP-independent variants of ShkA allowed us to more carefully investigate the role of ShkA in the temporal control of events during G1/S. Introduction of the *shkA^D369N^* allele into the rcdG^0^ strain restored SpmX protein levels, stalk biogenesis, DivJ localization to the incipient stalked pole, and normal cell morphology (Fig. 3a,b; Supplementary Fig. 6a). Of note, *shkA^D369N^* failed to restore G2-specific processes in the rcdG^0^ strain like assembly of the flagellum and type IV pili (Supplementary Fig. 6b,c). Restoration of cell morphology and DivJ localization was entirely dependent on SpmX (Fig. 3b), arguing that c-di-GMP and ShkA drive cell cycle progression and morphogenesis via *spmX* expression control. However, a strain harboring the *shkA^D369N^* allele prematurely produced SpmX already in newborn G1 cells (Fig. 3c). Accordingly, the *shkA^D369N^* mutant exited G1 prematurely, as indicated by the strong reduction of G1 cells (Fig. 3d). Likewise, cells failed to properly arrest in G1 after entry into the stationary phase (Fig. 3d,e). These results support a role of c-di-GMP, and the ShkA-TacA pathway, in the timely execution of the G1/S transition.

**Fig. 3:**
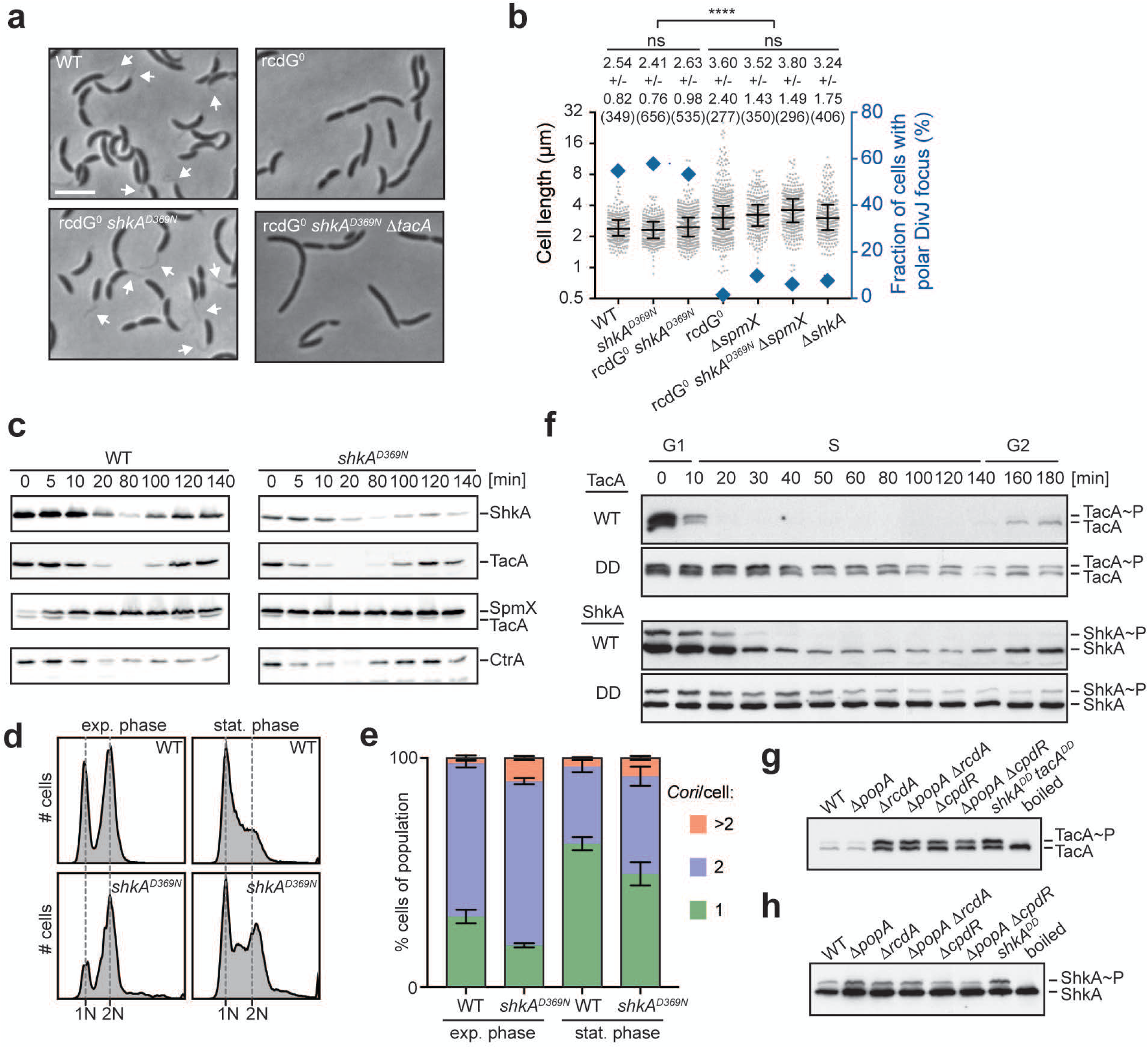
C-di-GMP-mediated activation and termination of the ShkA-TacA phosphorelay imposes precise temporal control during G1/S. **a**, Representative phase-contrast micrographs of indicated strains. Arrows point to stalks. The scale bar represents 4 µm. **b**, Quantification of cell length and polar DivJ localization (DivJ-mCherry) of indicated strains. Median values with interquartile ranges are shown in the graph and mean values and standard deviations are indicated above the graph. The number of cells analyzed is shown in brackets. **** indicates a P value of <0.0001; ns, not significant. **c**, Immunoblots of synchronized cultures of *C. crescentus* strains expressing *3xFLAG*-*shkA* or *3xFLAG*-*shkA^D369N^* from the native chromosomal locus were probed with anti-FLAG, anti-TacA, anti-SpmX and anti-CtrA antibodies. **d**, Analysis of chromosome content by flow cytometry of indicated strains in exponential or stationary phase after rifampicin treatment. **e**, Quantification of chromosome number of indicated strains. Shown are means and standard deviations (N=3). **f**, Phos-tag PAGE immunoblots of synchronized cultures of strains expressing chromosomally encoded *3xFLAG*-*tacA*, *3xFLAG*-*shkA, 3xFLAG*-*tacA^DD^* or *3xFLAG*-*shkA^DD^* alleles were probed with anti-FLAG antibody. **g**, Phos-tag PAGE immunoblots of mixed cultures of indicated mutant strains expressing *3xFLAG*-*tacA* probed with anti-FLAG antibodies. **h**, Phos-tag PAGE immunoblots of mixed cultures of indicated mutant strains expressing *3xFLAG*-*shkA* probed with anti-FLAG antibodies.

TacA was previously shown to be degraded by the ClpXP protease, a process that requires the adaptor proteins RcdA and CpdR and depends on the C-terminal Ala-Gly degradation motif of TacA ^15^. ShkA harbors the same C-terminal degradation signal and was also degraded upon S-phase entry about 10 minutes after the removal of TacA (Fig. 3f). When the C-terminal Ala-Gly motif was replaced by Asp residues (ShkA^DD^), ShkA and ShkA∼P were stabilized and prevailed throughout the cell cycle (Fig. 3f). Sequential degradation of TacA and ShkA may be explained by their differential requirements for protease adaptors. While TacA degradation by ClpXP depends on CpdR and RcdA ^15^, ShkA degradation also requires PopA, the third member of the adaptor hierarchy regulating ClpXP protease activity during the *C. crescentus* cell cycle (Fig. 3g,h). Because PopA needs to bind c-di-GMP to act as a protease adaptor ^16^, the ShkA-TacA pathway is confined to G1/S by the sequential c-di-GMP-dependent activation and c-di-GMP-mediated degradation of the ShkA kinase.

### The ShkA-TacA pathway limits gene expression to G1/S

To carefully assess the contribution of ShkA activation and degradation for the temporal control of *spmX,* the *spmX* promoter was fused to the fluorescent protein Dendra2. The photoconvertible properties of Dendra2 ^17^ allowed determining both ‘ON’ and ‘OFF’ kinetics of *spmX* promoter activity during the cell cycle (Fig. 4a; Supplementary Fig. 7a; for details, see Materials and Methods). These experiments revealed that *spmX* promoter activity peaks during G1/S roughly 15-30 minutes after passing through the predivisional stage (Fig. 4b). The *spmX* promoter was active in cells progressing through G1/S, but not in newborn ST progeny that re-enter S-phase immediately at the end of the asymmetric cell cycle (Fig. 4a,c; Supplementary Fig. 7a-c). Stage-specific expression of *spmX* required a combination of c-di-GMP oscillations during the cell cycle and TacA degradation: expression of combinations of *dgcZ*, a gene encoding a highly active DGC from *E. coli* ^18^, and of *tacA^DD^* (encoding stable TacA) and *shkA^D369N^* (encoding constitutive ShkA) alleles resulted in a gradual loss of G1/S-specific *spmX* expression (Fig. 4c; Supplementary Fig. 7c). These experiments provided direct evidence that the activity of the ShkA-TacA pathway is strictly limited to G1/S and SW progeny that need to passage through the G1 phase of the cell cycle.

**Fig. 4:**
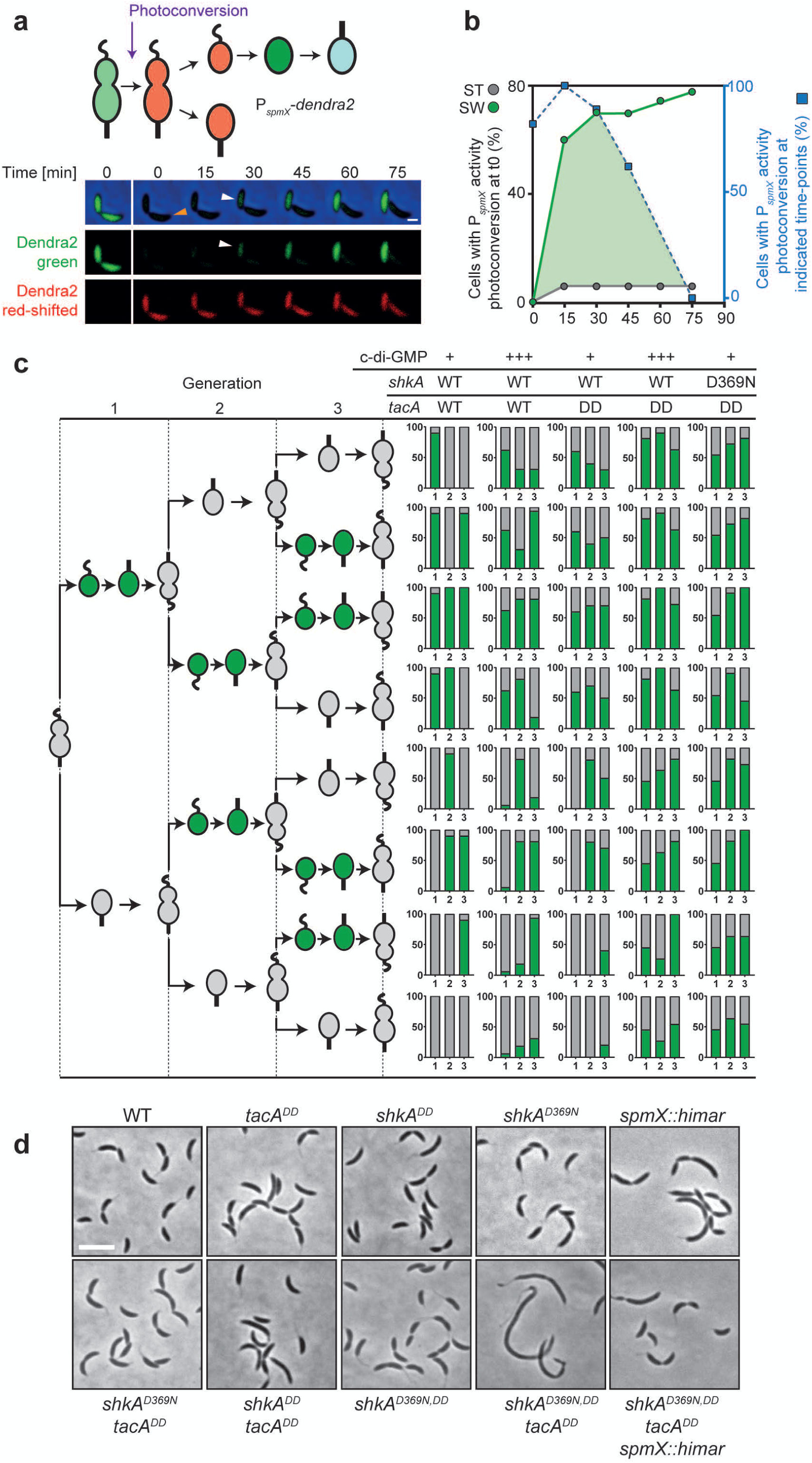
Oscillation of c-di-GMP and protein degradation limit TacA activity to G1/S. **a**, Time-lapse fluorescence microscopy of *C. crescentus* wild-type cells expressing *dendra2* from the *spmX* promoter. The time of photoconversion of Dendra2 (green to red) in the predivisional cell (PD) is indicated in the schematic of the *C. crescentus* cell cycle. A representative example of a dividing cell is shown below with separate green (FITC) and red (TRITC) channels and with the ST cell pole of the PD cell marked (orange arrow). The *spmX* promoter (green) is activated exclusively in the SW cell during G1/S transition (white arrows). The bar is 1 µm. **b**, ON and OFF kinetics of *spmX* promoter activity during G1/S in *Caulobacter* wild-type cells harboring the *spmX-dendra2* reporter plasmid. ON kinetics were determined as outlined in **a** with cells being photoconverted before division (0 min) and the fraction of ST (grey circles) and SW cells (green circles) with induced green fluorescence plotted over time. OFF kinetics were determined by the fraction of SW cells with induced green fluorescence after photoconversion at the indicated time points during the cell cycle (blue boxes). The overlap of the two curves (green area) defines the window of *spmX* promoter activity during the cell cycle. **c**, Activity of *spmX* promoter in lineages of individual *C. crescentus* cells of different strains through three consecutive generations as indicated by the schematic on the left. For each strain 10-16 late PD harboring the *spmX-dendra2* reporter were photoconverted roughly 15 minutes before cell division and followed by time-lapse microscopy through three cell division events. Right: The fraction of SW and ST offspring with active *spmX* promoter (green) was plotted over three generations with the x-axis representing generations 1-3. For experimental details and data analysis see Materials and Methods. C-di-GMP levels were manipulated by P*lac*-driven *dgcZ* with 0.1 mM IPTG. EV, empty vector control. **d**, Representative phase-contrast micrographs of strains carrying different *shkA* and *tacA* alleles encoding stabilized (*shkA^DD^* and *tacA^DD^*) or c-di-GMP-independent (*shkA^D369N^*) versions of the respective proteins. The scale bar represents 4 µm.

To investigate the importance of limiting the ShkA-TacA pathway to G1/S for accurate cell cycle progression, we examined the consequences of ShkA-TacA dysregulation using stable (TacA^DD^, ShkA^DD^), or constitutively active (ShkA^D369N^) variants or combinations thereof. All alleles increased overall *spmX* expression and showed additive effects when combined (Supplementary Fig. 8a). Cell division and cell morphology were normal in all strains that either retained TacA degradation or c-di-GMP-mediated ShkA activity control. However, when mutations that stabilize ShkA or TacA were combined with a mutation constitutively activating ShkA, strains showed strong cell division and morphology aberrations, effects that were strictly dependent on an intact copy of *spmX* (Fig. 4d; Supplementary Fig. 8b). Thus, dysregulation of the SpmX morphogen during G1/S leads to aberrant cell morphogenesis. Together, these experiments provide evidence that the activity of the ShkA-TacA pathway is strictly limited to G1/S and that dysregulation of this narrow temporal window of activity leads to severe cell cycle and morphological defects.

### The diguanylate cyclase PleD activates ShkA during G1/S

The above data support a model in which an upshift of c-di-GMP stimulates ShkA kinase activity thereby initiating the G1/S-specific genetic program (Fig. 5a). If so, the G1/S transition should be kick-started by one of the *C. crescentus* diguanylate cyclases. Screening a *spmX-lacZ* reporter strain for transposon insertions with reduced *lacZ* expression identified mutations in *pleD*. Accordingly, a *ΔpleD* deletion strain showed strongly reduced *spmX* expression (Fig. 5b). While a second diguanylate cyclase enzyme, DgcB, had a more modest effect, *spmX* promoter activity was almost completely abolished in a *ΔpleD ΔdgcB* double mutant, akin to a strain lacking c-di-GMP (Fig. 5b). Thus, PleD is the major diguanylate cyclase driving the G1/S-specific transcriptional program.

**Fig. 5:**
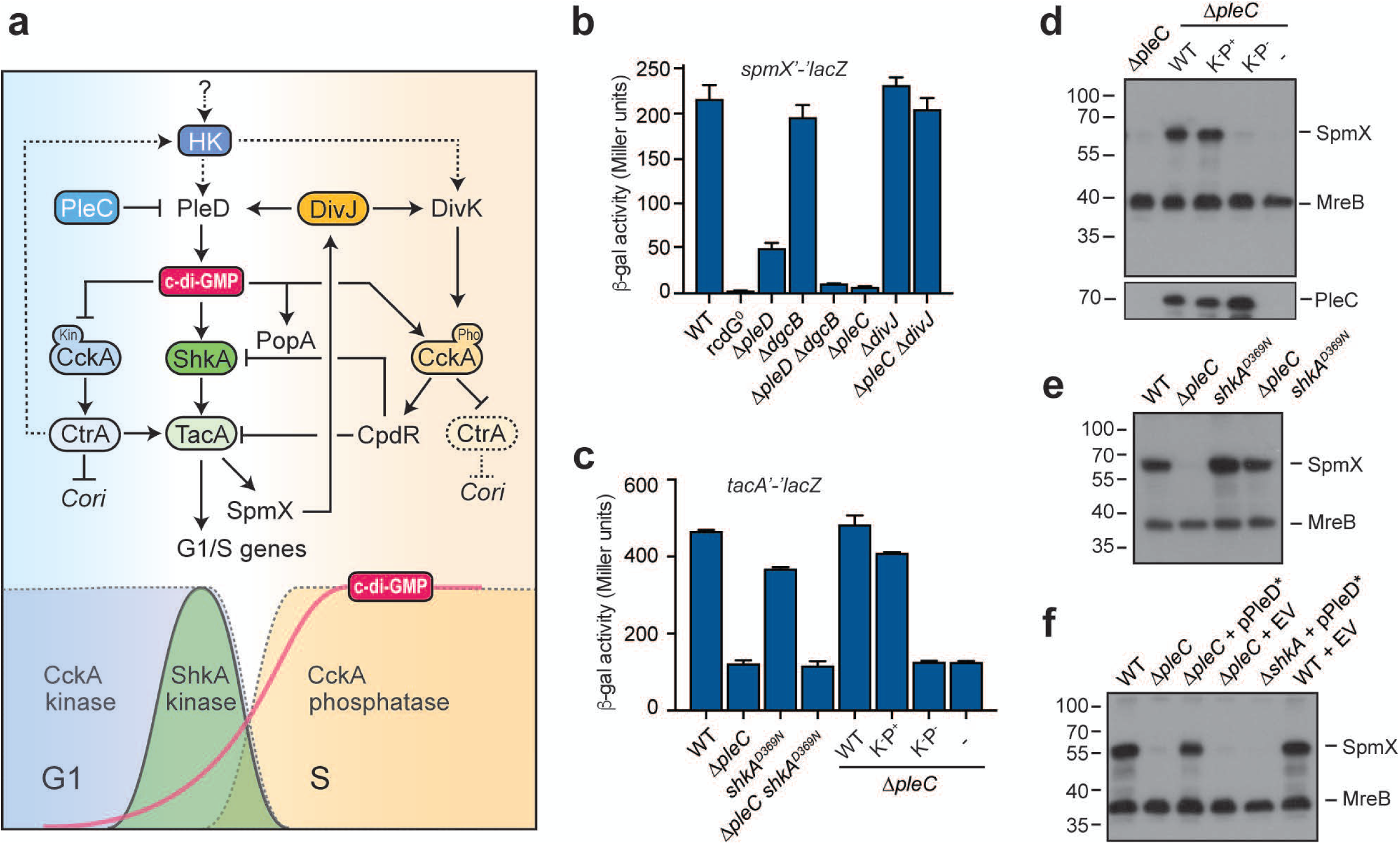
C-di-GMP-dependent ShkA activation requires PleC and PleD. **a**, Model of ShkA-TacA phosphorelay activation and its contribution to G1/S transition. The upper part shows the intercalated network of kinases (PleC, DivJ, ShkA, CckA) and response regulators (PleD, TacA, DivK) that contribute to G1/S transition. HK indicates a hypothetical histidine kinase that controls the activity of PleD and possibly DivK (stippled lines) and by that acts as kick-starter for G1 exit. HK expression is postulated to be CtrA-mediated (stippled line). The initial, HK-PleD-mediated increase of c-di-GMP activates the ShkA-TacA pathway, resulting in a SpmX-DivJ-PleD-mediated further boost of c-di-GMP concentration, which activates the CckA phosphatase and PopA. Note that ShkA has a 5-10-fold higher affinity for c-di-GMP compared to CckA and PopA. The block of replication initiation by binding of activated CtrA to the chromosomal origin of replication (*Cori*) is indicated. The lower part of the graph indicates time windows of CckA kinase and phosphatase as well as TacA kinase activity. The gradual increase of c-di-GMP during G1/S is indicated. **b**, Activity of the *spmX* promoter in strains harboring a *spmX’*-‘*lacZ* reporter fusion (pAK502-*spmX*). Shown are mean values and standard deviations (N=3). **c**, Activity of the *tacA* promoter in strains harboring a *tacA’-‘lacZ* fusion (pAK502-*tacA*). Strains with a chromosomal *pleC* deletion express wild-type and mutant *pleC* alleles from plasmid pQF under control of their native promoter. PleC(F778L): Kinase-/phosphatase+ (K-P+); PleC(T614R) kinase-/phosphatase-(K-P-); - indicates the empty vector control (plasmid pQF). Shown are mean values and standard deviations (N=3). **d**, Immunoblots of selected strains shown in panel **c** probed with anti-SpmX and anti-MreB antibodies (top) or anti-PleC antibodies (bottom). **e**, Immunoblots of selected strains shown in panel **c** probed with anti-SpmX and anti-MreB antibodies. **f**, Immunoblots of indicated strains probed with anti-SpmX and anti-MreB antibodies. pPleD* indicates a pMR20-based plasmid (pPA114-47) expressing the constitutively active, phosphorylation-independent PleD* allele. EV denotes the empty plasmid control (pMR20).

PleD activity is inversely regulated by the ST cell-specific kinase DivJ and the SW cell-specific phosphatase PleC (Fig. 5a) ^8^. However, PleC but not DivJ, was required for *spmX* expression, and *spmX* expression was restored to normal levels in a strain lacking both PleC and DivJ (Fig. 5b). This indicated that neither DivJ nor PleC is responsible for the initial activation of PleD and for ShkA stimulation, and that the role of PleC is likely indirect. PleC phosphatase was previously shown to reduce DivK phosphorylation leading to the activation of CtrA. Because *tacA* is a direct target of CtrA, it was proposed that in a *ΔpleC* mutant, *spmX* expression is impaired because TacA fails to accumulate ^12^. Expression of *tacA* and *spmX* indeed required the PleC phosphatase (Fig. 5c,d). However, when the *pleC* deletion was combined with the constitutive *shkA^D369N^* allele, SpmX protein levels and polar localization of DivJ, but not *tacA* expression, were fully restored (Fig. 5c,e; Supplementary Fig. 9), arguing that TacA levels are not the limiting factor in cells lacking PleC. Rather, ShkA activity and its stimulation by c-di-GMP are switched off in the *ΔpleC* mutant. Indeed, a constitutively active variant of PleD, PleD* ^14^, restored SpmX protein levels in the *ΔpleC* mutant, an effect that was entirely dependent on ShkA (Fig. 5f). These results suggest that c-di-GMP levels are limiting in a *ΔpleC* mutant. In line with this, the *ΔpleC* mutant showed significantly reduced c-di-GMP levels as compared to the isogenic *pleC*^+^ strain (9.5 ± 1.3 µM vs. 16.3 ± 0.5 µM; N=3).

The finding that the ShkA-TacA pathway is OFF in the *ΔpleC* mutant because c-di-GMP concentrations are limiting, together with the observation that PleD serves as the main c-di-GMP donor for ShkA activation, argues for the existence of an as yet unidentified PleD kinase, the expression of which will likely depend on the PleC-CckA-CtrA cascade. We speculate that activation of this kinase and its downstream target PleD represents a key event in the decision of *C. crescentus* to exit G1 (Fig. 5a).

## DISCUSSION

*C. crescentus* SW cells are born with low levels of c-di-GMP ^11,19^. This is imposed by two cell type-specific regulators, the phosphodiesterases PdeA ^20^ and the phosphatase PleC, which maintains the diguanylate cyclase PleD in its inactive, unphosphorylated form ^8^ (Fig. 5a). During the G1/S transition, PdeA is proteolytically removed ^20^ and PleD is activated by phosphorylation. This results in a gradual increase of c-di-GMP ^11,19^, which leads to a series of accurately timed events prompting exit from G1, cell morphogenesis and entry into S-phase. First, ShkA is allosterically activated resulting in TacA phosphorylation and the expression of a large group of G1/S-specific genes that orchestrate the morphological restructuring of the motile SW cells into sessile ST cells ^13,21^. We presume that at this stage, c-di-GMP levels are high enough to activate ShkA-TacA but may not have reached the peak levels needed to trigger the CckA cell cycle switch and S-phase entry ^8,9^. This leads to the execution of the morphogenetic program before cells commit to chromosome replication and division. The next step is then catalyzed by the expression of one of the G1/S-specific proteins, the morphogen SpmX, which is responsible for the polar sequestration and activation of DivJ ^12^. This, in turn, leads to the production of more c-di-GMP via reinforced activation of PleD and, together with DivJ-mediated phosphorylation of DivK, switches CckA into a phosphatase and ultimately licenses replication initiation ^7–9^ (Fig. 5a). ShkA binds c-di-GMP with 5-10-fold higher affinity than CckA ^9,16^ explaining how the ShkA-TacA pathway and the CckA switch can be sequentially activated. Thus, at least four kinases form a hierarchical cascade (HK→ShkA→DivJ→CckA) that is responsible for the accurate temporal control of events during G1/S. The activity and timing of this cascade is coordinated by the second messenger c-di-GMP, a stepwise increase of which enforces consecutive cell cycle steps by modulating the activity of ShkA and CckA, respectively (Fig. 5a).

By contributing to the CckA phosphatase switch via SpmX and DivJ, the ShkA-TacA pathway initiates its own termination. The CckA phosphatase activates a protease adaptor cascade that includes CpdR and PopA ^15^ and that leads to the consecutive degradation of TacA and ShkA by the ClpXP protease (Fig. 5a). This negative feedback constitutes an intrinsic, self-sustained timer that shuts down ShkA-TacA activity as soon as c-di-GMP has reached peak levels required to activate the CckA phosphatase and the PopA protease adaptor, thereby irreversibly committing cells to S-phase. Because TacA not only controls genes involved in cell cycle progression but also regulates morphological restructuring of the motile SW cells into sessile ST cells ^13,21^, accurate temporal control of this pathway may secure the tight coordination between replicative and behavioral processes.

Our results show that the diguanylate cyclase PleD is largely responsible for the c-di-GMP upshift during G1-S transition. We postulate that the initial event leading to PleD activation during G1/S must be executed by a kinase other than DivJ or PleC, and that DivJ is part of a positive feedback loop that reinforces PleD activity upon S phase entry (Fig. 5a). Although the nature of this kinase is currently unknown, we speculate that its expression or activity is CtrA-dependent and that it plays a key role in orchestrating exit from G1 as it may not only serve to activate PleD and provide the initial boost of c-di-GMP but may also contribute to the activation of DivK, a factor required for the CckA kinase/phosphatase switch. The essential nature of DivK but not of DivJ, its only known activating kinase, argues for regulatory redundancy in the upstream components required to boost DivK phosphorylation during G1/S (Fig. 5a).

We postulate that in its default state, the ShkA kinase is inhibited by the C-terminal REC2 domain and that c-di-GMP binding liberates the kinase by interfering with this off-state conformation. A ShkA variant lacking the REC2 domain is active without c-di-GMP ^13^. Similarly, mutations of residues important for REC2 function lead to constitutive, c-di-GMP-independent autokinase activity. Thus, the conserved DDR motif in the REC1-REC2 linker likely serves to lock ShkA in the inactive state when no c-di-GMP is present. REC2 may be closely tethered to REC1 in the inactive state through an interaction of the DDR linker motif with REC1, a conformation that may prevent the productive interaction of the catalytic CA with the DHp domain. We hypothesize that binding of c-di-GMP to REC1 interferes with this tethering, thereby liberating REC2 and facilitating the productive interaction between CA and DHp for autophosphorylation and eventually for phosphotransfer between DHp and REC2. This model is strongly supported by an accompanying structural analysis of ShkA ^22^.

These findings demonstrate that degenerate REC domains, also called pseudo-receiver domains, can function as docking sites for small regulatory molecules. Hybrid histidine kinases with pseudo-receiver domains located between the CA and the phosphorylated receiver domain are widespread in bacteria and include the well-studied virulence factors of the GacS/BarA family ^23,24^ or the global stress regulator RcsC ^25,26^ (Supplementary Fig. 5). For instance, the pseudo-receiver of RcsC can only be recognized by structural comparison ^25^, arguing that primary structure-based searches largely underestimate the actual number of these modules. It is possible that pseudo-receiver domains generally serve as binding sites for metabolites or small signaling molecule thereby modulating kinase or phosphatase activity of such key bacterial regulators. In line with this, prominent examples of kinases harboring pseudo-receiver domains like GacS in *P. aeruginosa* or BarA and RcsC in *E. coli* are part of complex signaling cascades that globally regulate bacterial physiology and behavior. Careful scrutiny of the exact function of these domains will be essential to improve our understanding of such important regulatory nodes in bacteria.

## AUTHOR CONTRIBUTION

Conceptualization, A.K., A.M.H., C.vA., B.D., T.S. and U.J.; Methodology, A.K., A.M.H., C.vA., J.N., R.B., S.H. and U.J.; Formal Analysis, A.K., A.M.H., C.vA., R.B., S.H. and U.J.; Investigation, A.K., A.M.H., C.vA., J.N., R.B., S.H. and U.J.; Resources, B.D and T.S.; Writing – Original Draft, A.K., A.M.H., C.vA., R.B., S.H. and U.J. with contributions from all other authors; Funding Acquisition, A.M.H., S.H., T.S. and U.J.

## ACKNOWLEDGEMENTS

We thank Tim Sharpe (Biophysics Facility, Biozentrum, University of Basel), Alexander Schmidt (Proteomics Core Facility, Biozentrum, University of Basel), Janine Bögli and Stella Stefanova (FACS Core Facility, Biozentrum, University of Basel) for technical guidance, Christian Lori and Benoit-Joseph Laventie for preparation of c-di-GMP, Fabienne Hamburger for cloning and strain construction and Life Science Editors for editing assistance. We thank Patrick Viollier, Lucy Shapiro, Justine Collier, Joseph Chen, Matthew Malvey, Régis Hallez and Julia Vorholt for providing antibodies, plasmids and strains. This work was supported by the European Research Council (ERC) Advanced Research Grant (3222809) and the Swiss National Science Foundation (310030B_147090) to U.J. A.M.H. is a Marie Heim-Vögtlin fellow of the Swiss National Science Foundation (PMPDP3_171306) and was supported by a Human Frontier Science Program (HFSP) Long-Term Postdoctoral Fellowship (LT000779/2013) and a University of Basel Stay-on-track grant. The authors declare no competing interests.

## Accession code

The sequence-specific backbone resonance assignment of the ShkARec1 domain has been submitted to the Biological Magnetic Resonance Data Bank under the following accession code: 27768.

## SUPPLEMENTARY FIG. LEGENDS

**Supplementary Fig. 1.**
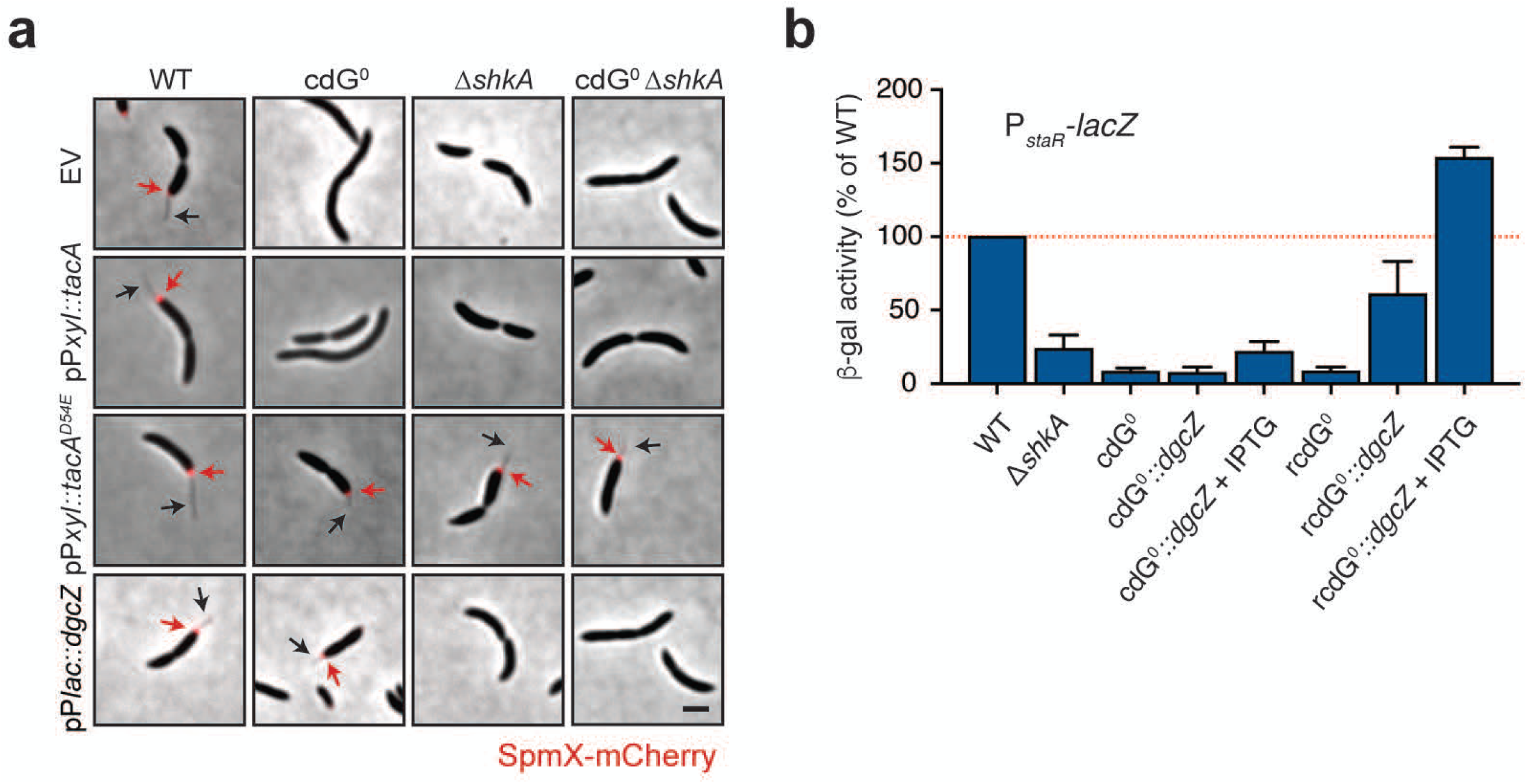
**a**, Micrographs of strains expressing a chromosomal *spmX-mCherry* fusion and plasmid-driven *tacA*, *tacA^D54E^* or the heterologous diguanylate cyclase *dgcZ*. EV, empty vector control. Stalks (black arrows) and SpmX-mCherry foci (red arrows) are marked. **b**, β-Galactosidase activities of indicated strains harboring a P*spmX*-*lacZ* transcriptional fusion (plasmid pRKlac290-*staR*). “+ IPTG” indicates induction of *dgcZ* expression from P*lac* by addition of 200 µM IPTG. Shown are means and standard deviations (N>3).

**Supplementary Fig. 2.**
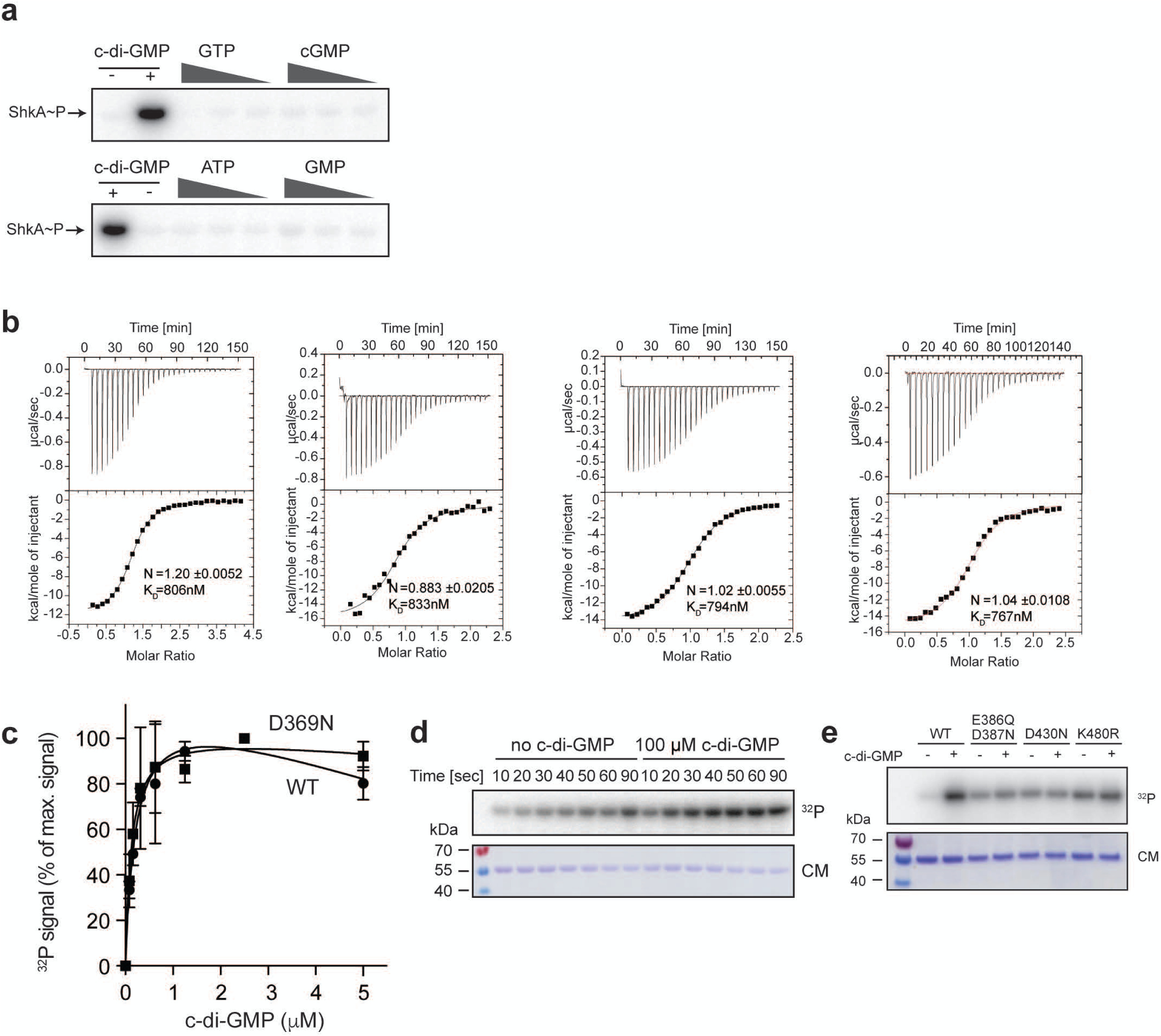
**a**, *In vitro* phosphorylation assays with purified ShkA in the presence of different concentrations of nucleotides (1 mM, 100 µM, 10 µM). Positive and negative controls contained 100 µM c-di-GMP and ddH2O, respectively. Reactions were initiated by addition of 500 µM radiolabeled ATP and allowed to proceed for 15 minutes at room temperature. **b**, Isothermal titration calorimetry measurements show a direct interaction of ShkA with c-di-GMP. Four independent experiments are shown. **c**, Quantified autoradiographs of purified ShkA and the ShkA^D369N^ variant (0.5 µM) UV-crosslinked with increasing concentrations of [^32^P]c-di-GMP. Shown are mean values and standard deviations (N=2). **d**, Time course of ShkA^D369N^ autophosphorylation at 4°C with or without c-di-GMP. Top: autoradiograph; bottom: Coomassie stain of the same gel. **e**, *In vitro* autophosphorylation assays of wild-type ShkA and indicated mutant variants with (10 μM) or without c-di-GMP. Autophosphorylation was allowed to proceed for 5 minutes at room temperature. Top: autoradiograph; bottom: Coomassie stain of the same gel. Residues E386 and D387 are responsible for magnesium binding of REC2, D430 is the residue accepting the phosphoryl group and K480 plays a role in stabilization of phosphorylated D430.

**Supplementary Fig. 3.**
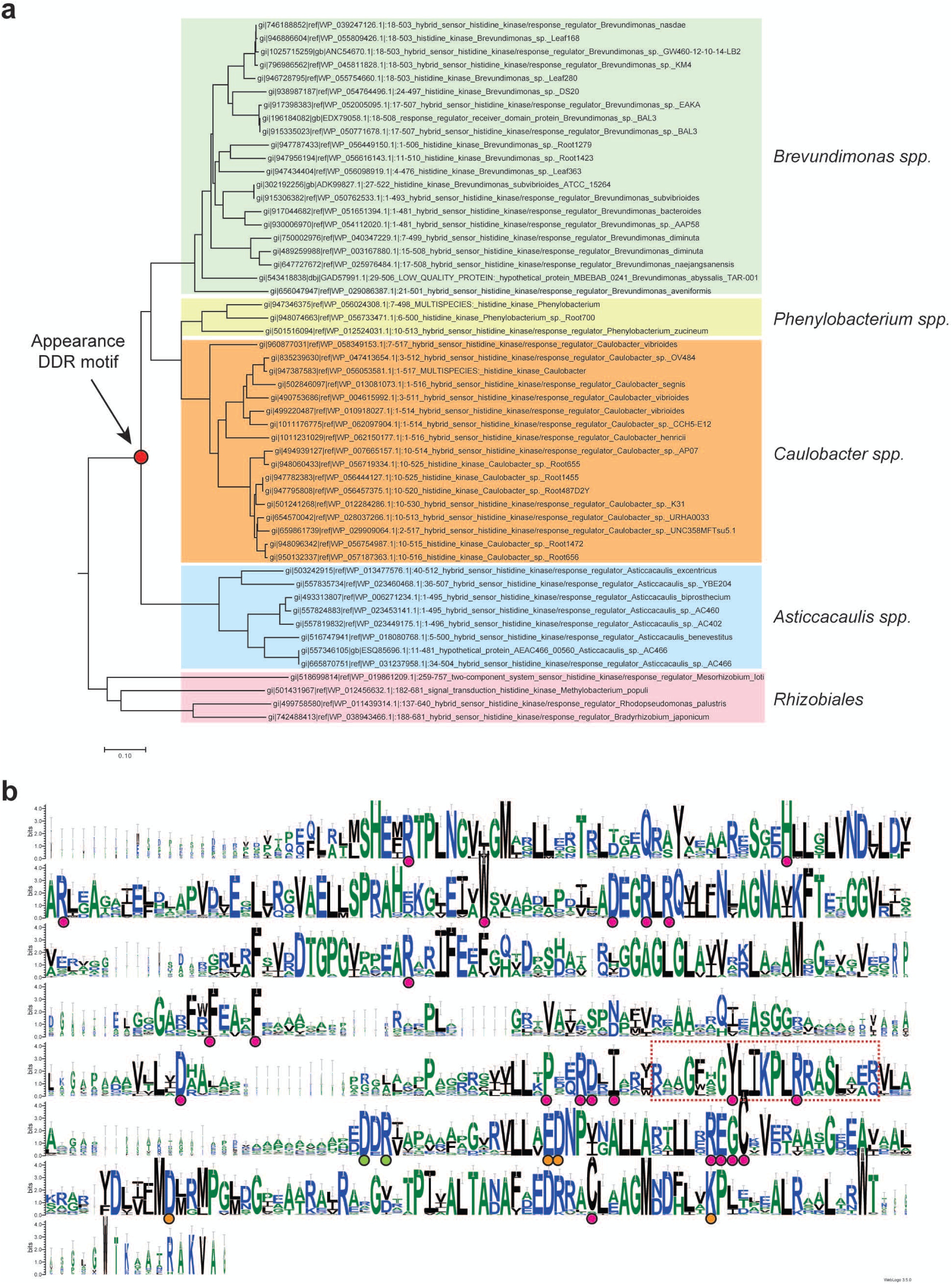
**a**, Phylogenetic tree of select ShkA orthologs created with Geneious Tree Builder using default parameters. **b**, Weblogo ^27^ based on an alignment of ShkA orthologs within the *Caulobacteraceae* shown in **a**. Circles below amino acid residues indicate residues that were mutated and tested for an effect on ShkA activity in *in vivo* (violet), were isolated as mutations that render ShkA activity c-di-GMP-independent (green), or were tested *in vitro* for their role in ShkA autoinhibition (orange). Conserved residues around the c-di-GMP binding site (see Fig. 2j,k) are highlighted by a box.

**Supplementary Fig. 4.**
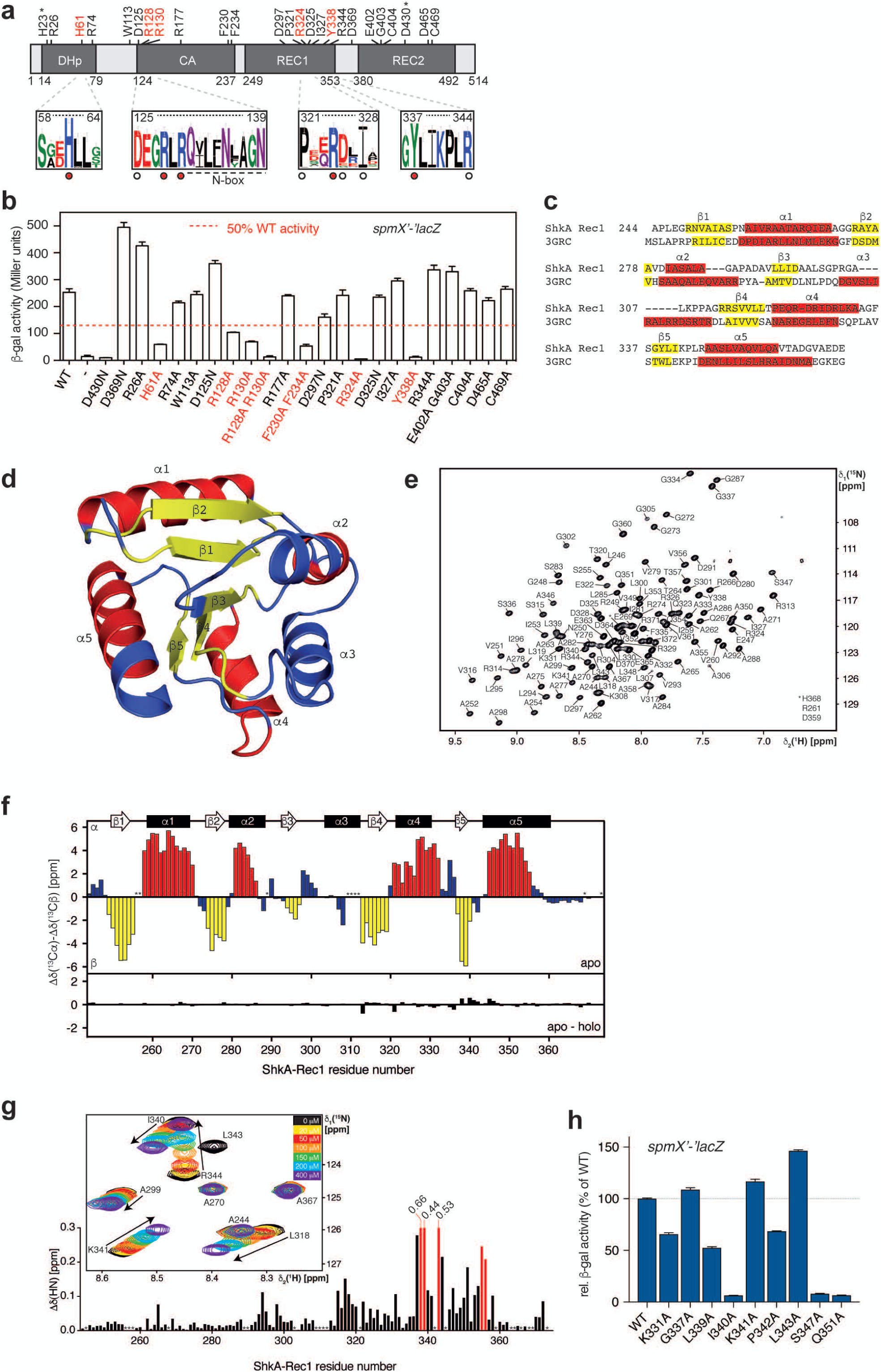
**a**, ShkA domain architecture (top) with domain boundaries indicated by amino acid numbers. Residues that were mutagenized and tested *in vivo* (see Supplementary Fig. 5) are indicated above the graph. Mutants that are impaired in c-di-GMP-dependent ShkA autophosphorylation are highlighted in red. These residues are also indicated by red dots below in Weblogos based on alignments of ShkA orthologs from different alphaproteobacteria (see Fig. S3). Residues that were mutagenized but did not affect ShkA activity *in vivo* are indicated by a white circle. The conserved phospho-acceptor His and Asp residues of the DHp and REC2 domain, respectively, are indicated by asterisks. **b**, β-Galactosidase activities of strain UJ9691 (NA1000 Δ*shkA* Δ*lacA*::Ω/pAK502-*spmX*) expressing indicated *shkA* alleles *in trans* from plasmid pQF. Note that no inducer (cumate) was present since leaky expression already restored the *shkA* null mutant phenotype. Shown are means and standard deviations (N=3). **c**, Profile-profile-based alignment of the REC domains of ShkAREC1 and sensor protein JS666 from *Polaromonas sp.* (PDB 3GRC) carried out with HHpred ^28^. Secondary structural elements (β-sheet, yellow; helix, red) are indicated as determined by NMR secondary chemical shifts for ShkAREC1 in solution (E) and by crystallographic data for PDB 3GRC (C). **d**, Published crystal structure of sensor protein JS666 from *Polaromonas sp.* (PDB 3GRC). The location of secondary structural elements (β-sheet, yellow; helix, red) of ShkAREC1 in solution are plotted onto the structure. **e**, 2D [^15^N,^1^H]-HSQC spectrum of 0.95 mM ShkAREC1 recorded at 25°C in 25 mM Tris pH 7.2 with 50 mM KCl and 2 mM MgSO_4_ in 95%/5% H2O/D2O. The sequence-specific resonance assignments are indicated. **f**, Sequence-specific secondary backbone ^13^C chemical shifts are plotted against the ShkA residue number, smoothed using a 1:3:1 weighting (top). Consecutive stretches with positive and negative values indicate α-helical (red bars) and β-strand (yellow bars) secondary structure, respectively. Asterisks indicate unassigned residues. Secondary chemical shift difference between apo and c-di-GMP-bound ShkAREC1 (bottom). **g**, Chemical shift perturbation of ShkAREC1 backbone amide moieties upon c-di-GMP binding. Combined chemical shift changes of amide moieties, Δδ(HN), are plotted against the residue number. The red bars indicate resonances that experience intermediate chemical exchange upon c-di-GMP binding. Asterisks indicate unassigned residues. Inset, region of a 2D [^15^N,^1^H]-HSQC spectrum from a titration of c-di-GMP to ShkAREC1 at 25°C in 25 mM Tris pH 7.2 with 50 mM KCl and 2 mM MgSO4 in 95%/5% H2O/D2O. **h**, β-Galactosidase activities of strain UJ9691 (NA1000 Δ*shkA* Δ*lacA*::Ω/pAK502-*spmX*) expressing indicated *shkA* alleles *in trans* from plasmid pQF. Note that no inducer (cumate) was present since leaky expression already restored the *shkA* null mutant phenotype. Values were normalized to the wild-type control assayed in parallel to the mutant alleles. Shown are means and standard deviations (N=2).

**Supplementary Fig. 5.**
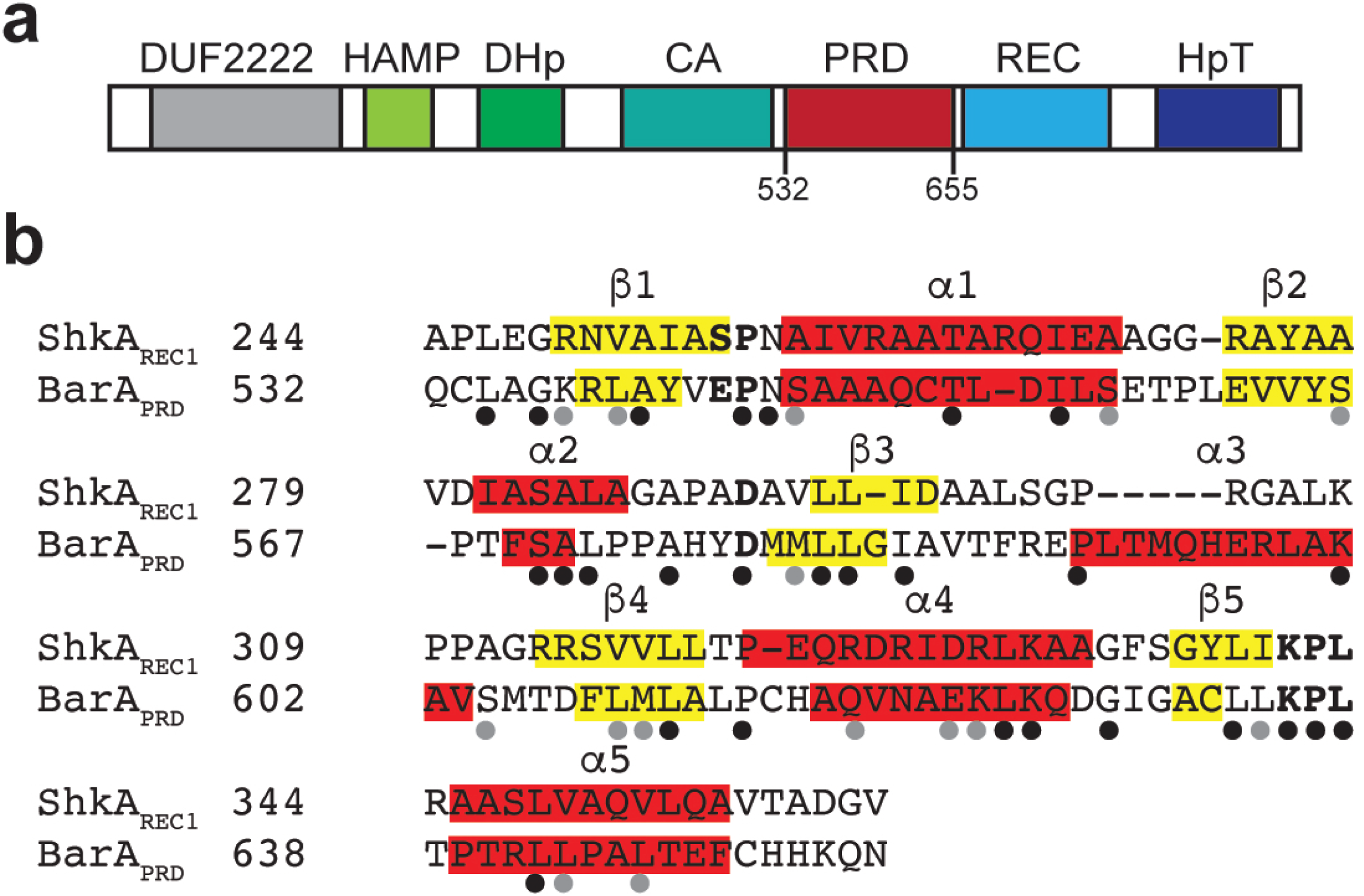
**a**, Domain architecture of *E. coli* K-12 BarA drawn approximately to scale. Domains were predicted using the SMART web server ^29^. The pseudo-receiver domain (PRD) is not predicted by SMART and was manually annotated (residues 532-655) based on the alignment shown in panel b. **b**, Sequence alignment of ShkAREC1 and BarA_PRD_ generated with the Geneious alignment tool. Secondary structures (yellow, *β*-sheets; red, *α*-helices) for ShkA_REC1_ and BarA_PRD_ are based on NMR data and HHPred predictions, respectively. Residues that are strictly conserved in prototypical REC domains are in bold and identical and similar residues between BarA_PRD_ and ShkA_REC_ are indicated by black and grey dots, respectively, below the sequence alignment. Note that a residue corresponding to the magnesium-binding pocket of canonical REC domains is degenerated in BarA (proline at position 591 instead of an acidic residue found in enzymatically active REC domains).

**Supplementary Fig. 6.**
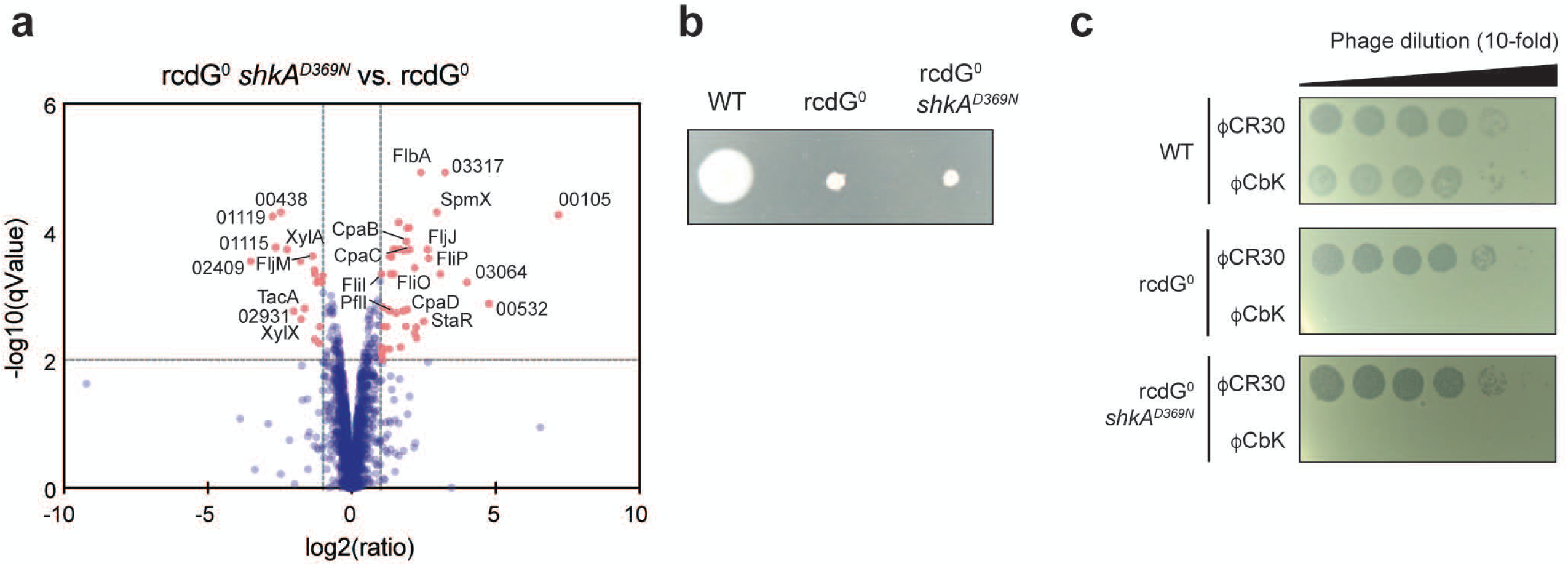
**a**, Volcano plot comparing the proteomes of the rcdG^0^ strain and its derivative carrying the s*hkA^D369N^* allele (strains SöA764 and UJ9619). Selected proteins that are differentially abundant in the two strains with high confidence and fold-change are labelled by their NA1000 CCNA number or their annotated gene product. **b**, Motility of indicated strains on semi-solid agar. Plates were incubated 3 days at 30°C. **c**, Sensitivity of indicated strains toward phage ΦCbK or ΦCR30 infection. Plates were incubated 24 hours at 30°C.

**Supplementary Fig. 7.**
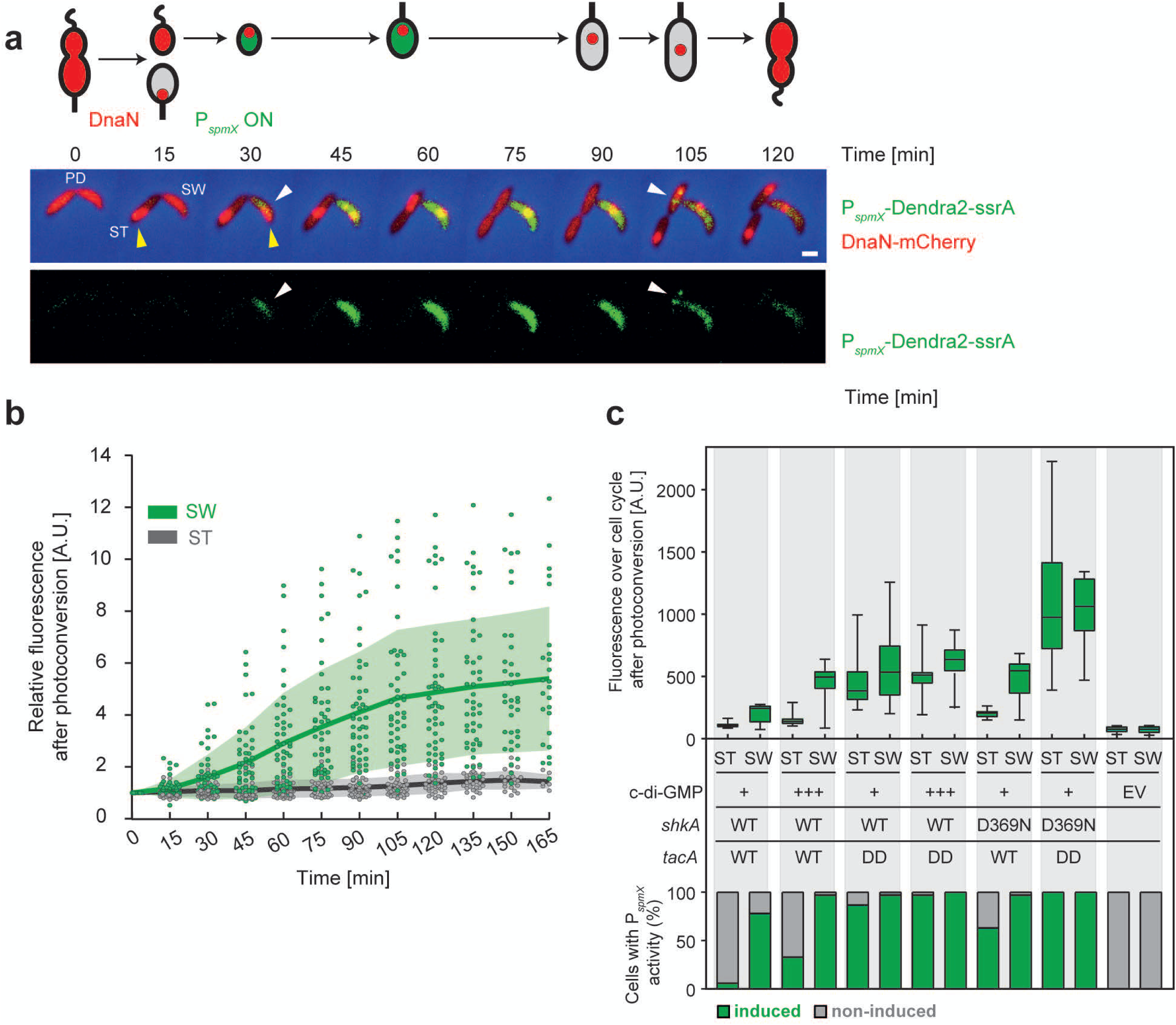
**a**, Representative example of a dividing *Caulobacter* wild-type cell harboring a transcriptional *spmX*::*Dendra2-ssrA* (green) reporter and a chromosomal *dnaN*-*mCherry* protein fusion (red). Transient localization of DnaN to the replisome at the incipient stalked pole (yellow arrow) indicates the start of S-phase in individual cells. A schematic (top) and fluorescence microscopy images (bottom) of dividing cells are shown at the indicated time points. Note that the *spmX* promoter is activated (Dendra2-ssrA, white arrows) prior to cells entering S-phase. The *ssrA*-tag facilitates proteolytic degradation of the fluorescent protein. The bar is 1 µm. **b**, Fluorescence of newborn wild-type SW (green dots) and ST progeny (grey dots) carrying P*spmX-dendra2* after photoconversion in their predivisional ancestors. The fluorescence intensity of individual cells is plotted as arbitrary units (A.U.) (N=50). Mean fluorescence (SW: green line; ST: grey line) and standard deviations (SW: green zone, ST: grey zone) are indicated. **c**, *spmX* promoter activity in newborn SW and ST cells measured by time-lapse microscopy of *C. crescentus* wild type and mutants harboring the *spmX-dendra2* reporter plasmid. For each strain, 30-50 late PD cells were photoconverted 15 minutes before cell division. Normalised fluorescence of individual SW and ST offspring over the next cell cycle was plotted as arbitrary units (A.U.) in quartiles in box-and-whiskers plots, the median indicated as black line and the whiskers extending from min to max values. The fraction of daughter cells with induced *spmX* promoter activity (green) is plotted in the bar chart below. For experimental details and data analysis see Materials and Methods. C-di-GMP levels were manipulated by P*lac*-driven *dgcZ* with 0.1 mM IPTG. EV, empty vector control.

**Supplementary Fig. 8.**
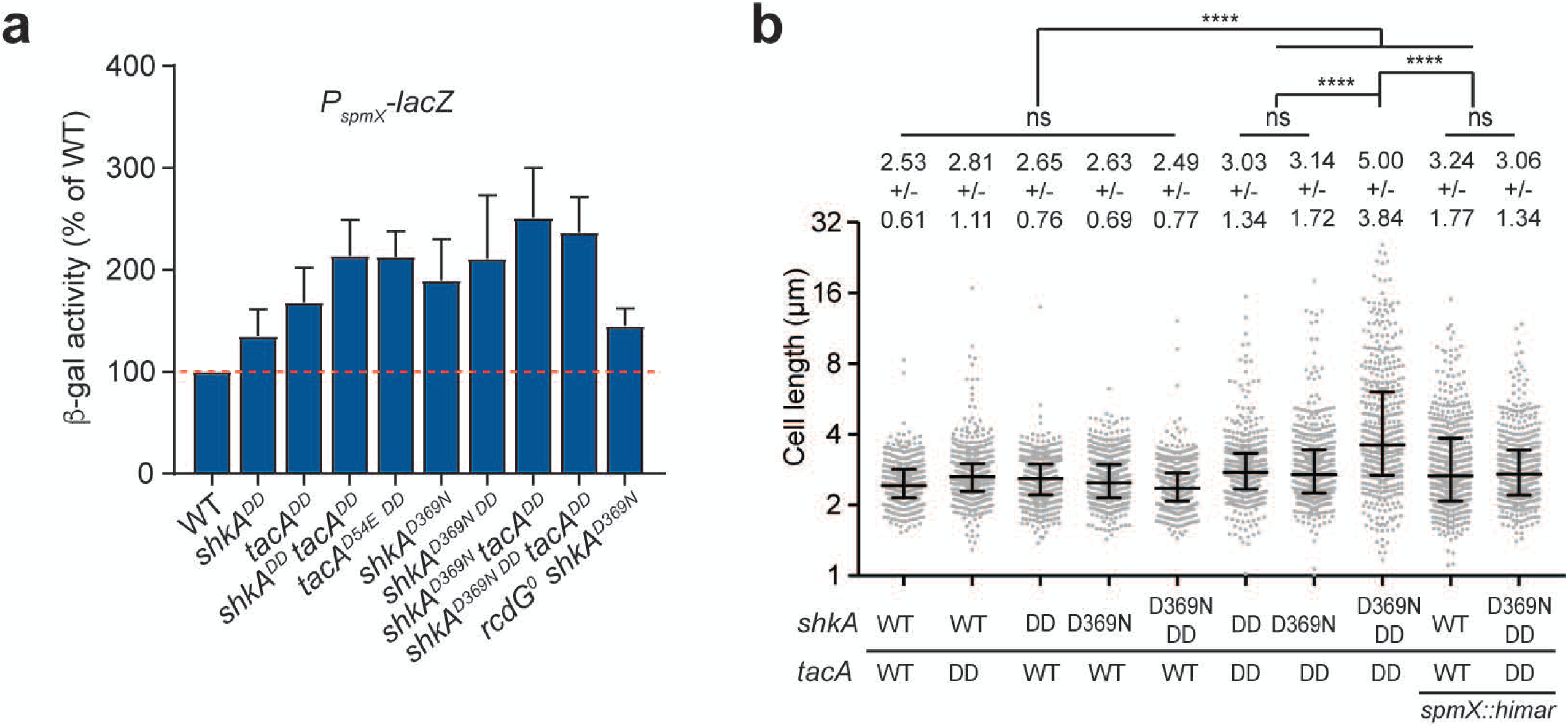
**a**, The expression of *spmX* can be increased by stabilizing proteins in the ShkA-TacA pathway as well as expressing constitutively active ShkA^D369N^. β-galactosidase assays were performed with strains carrying the pRKlac290-*spmX* reporter plasmid. Promoter activity of *spmX* was measured in wild-type and mutant strains and normalized to wild-type levels. Data represent means and standard deviations (N>3). **b**, Quantification of cell length of strains with mutations in *shkA* or *tacA* that block protein degradation (DD) or with constitutive c-di-GMP-independent ShkA activity (D369N). Median values with interquartile ranges are shown and the means and standard deviations are indicated above the graph. 480-489 cells were analyzed for each strain. *, **, *** and **** indicate P values of < 0.1, <0.01, <0.001 and <0.0001, respectively; ns, not significant.

**Supplementary Fig. 9.**
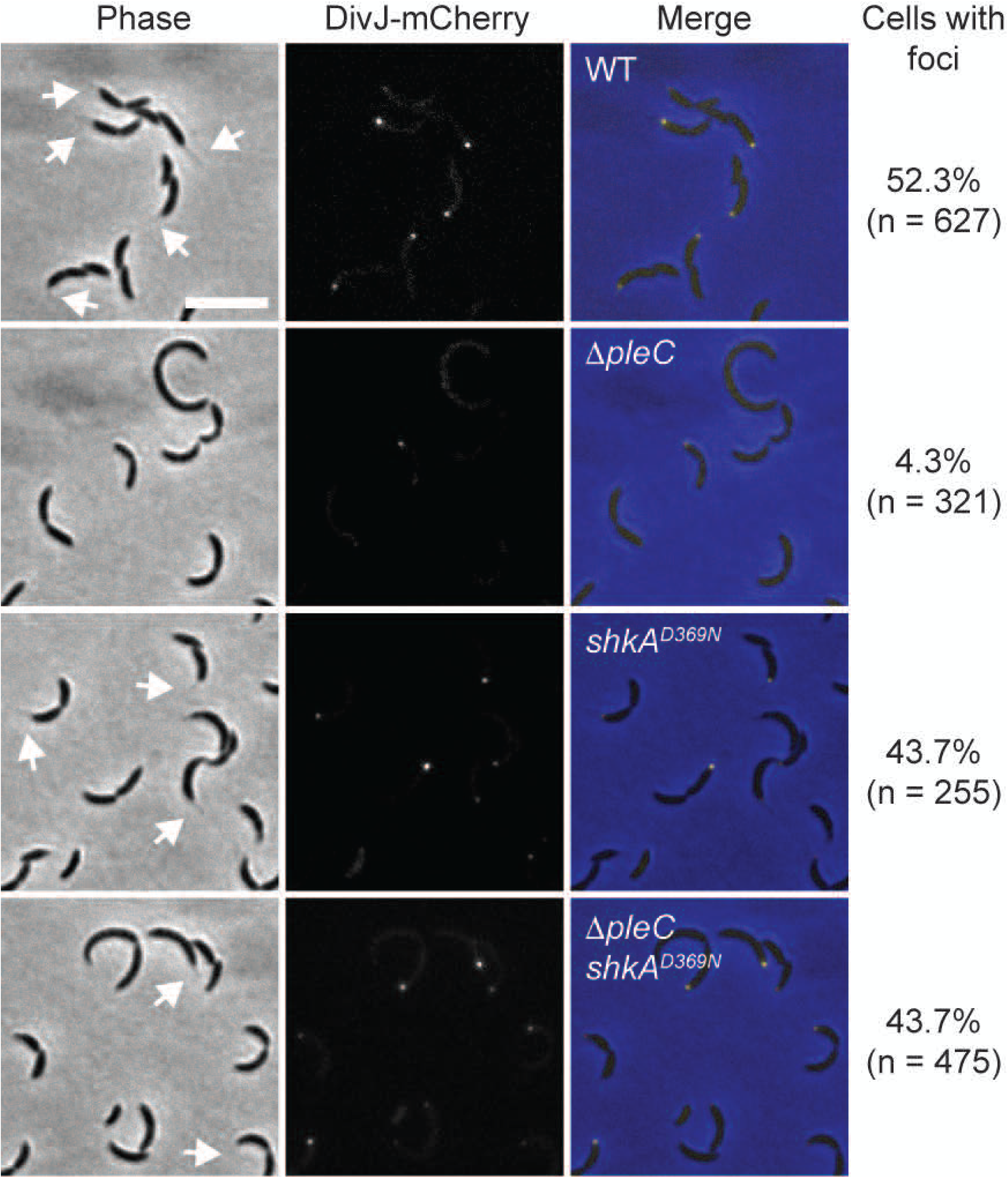
Stalk formation and polar localization of a chromosomally encoded DivJ-mCherry fusion was followed in wild-type, *ΔpleC, shkA^D369N^* and *ΔpleC shkA^D369N^* strains. White arrows in phase contrast imaged highlight stalks. The scale bar represents 4 µm.

## MATERIALS AND METHODS

### Growth conditions

*Caulobacter crescentus* was grown in PYE (0.2% [w/v] bacto peptone, 0.1% [w/v] yeast extract, 0.8 mM MgSO_4_, 0.5 mM CaCl_2_) or defined M2G (12.2 mM Na_2_HPO_4_, 7.8 mM KH_2_PO_4_, 9.3 mM NH_4_Cl, 0.5 mM MgSO_4_, 0.5 mM CaCl_2_, 20 uM FeSO_4_, 0.2% [w/v] D-glucose) medium at 30°C. Standard plates contained 1.5% [w/v] agar. Motility was assayed on PYE plates containing 0.3% agar. *Escherichia coli* DH5α, DH10B or TOP10 were used for cloning and were routinely cultivated in LB-Miller at 37°C. When appropriate, media were supplemented with antibiotics at the following concentrations unless stated otherwise (liquid/solid media for *C. crescentus*; liquid/solid media for *E. coli*; in µg/ml): kanamycin (5/20; 30/50), oxytetracycline (2.5/5; 12.5/12.5), chloramphenicol (1/2; 20/30), gentamycin (1/5; 20/20), spectinomycin (25/50; -/-), streptomycin (5/5; -/-), ampicillin (-/-; 100/100), nalidixic acid (15/20; -/-). Isopropy-β-D-thiogalactopyranoside (IPTG), 4-hydroxy-3-methoxybenzoic acid (vanillate) and xylose stocks were prepared in ddH2O at concentrations of 1 M, 200 mM and 20% [w/v], respectively. 4-isopropylbenzoic acid (cumate) stocks were prepared in 100% ethanol.

### Strains and plasmids

Plasmids are listed in Table S1. Oligonucleotides used for plasmid construction were purchased from Sigma and are listed in Table S2, and plasmid construction details are described in the Supplementary Methods. Plasmid were delivered into *C. crescentus* by electroporation or triparental mating using LS980 as helper strain. φCR30 was used for generalized transduction ^30^. Strains are listed in Table S3. In general, *C. crescentus* mutant strains were constructed by a two-step homologous recombination selection/counterselection procedure using pNPTS138. Alternatively, pNPTS138-based plasmids used for construction of a specific mutant strain were first integrated in this strain and then the mutation was moved in a new background using φCR30-mediated generalized transduction, followed by resolution of the merodiploid by counterselection on PYE plates containing 0.3 % [w/v] sucrose.

### Synchronization

Strains were grown overnight in 5 ml PYE supplemented with appropriate antibiotics and inducers in a roller drum at 30°C. The next day, cultures were diluted 20-fold in 25 ml M2G and allowed to grow on a rocking shaker at 30°C until they reached an OD_660_ of 0.4 to 0.6. These cultures were then subcultured in 300 ml M2G such that the cultures reached an OD_660_ of 0.6 after overnight incubation on a rocking shaker (160 rpm) at 30°C. Cells were harvested (5000 g, 10 min, 4°C), the supernatant was aspirated, cells were resuspended in 40 ml cold phosphate buffer (12.3 mM Na2HPO4, 7.8 mM KH2PO4) for experiments shown in Fig. 4b or in cold M2G for experiments shown in Fig. 4a and kept at 4°C for the rest of the procedure. 20 ml cold Ludox were added and the suspension mixed thoroughly before being transferred to pre-cooled glass Corex tubes and spun to separate the swarmer from the stalked cells (9780 g, 35 min, 4°C). The top layer of cells was aspirated, the swarmer band moved to a clean pre-cooled Corex tube and washed twice with 30 ml of either cold phosphate buffer or M2G (see above) by centrifugation (8000 g, 10 min, 4°C). Cells were released to a flask with pre-warmed M2G at an OD_660_ of approximately 0.3 and incubated in a water bath (30°C, 150 rpm). Samples were taken at indicated time-points in pre-cooled tubes and harvested at max. speed using a cooled table-top centrifuge. The pellets were immediately snap-frozen in liquid nitrogen and stored at −20 °C until further processing.

### Phos-tag PAGE

Samples from mixed cultures grown in PYE were taken in exponential phase, and sampling of synchronized cultures is described above. An equivalent of 0.5 ml cell culture at an OD_660_ of 0.5 was harvested by centrifugation in a table-top centrifuge at max speed and the pellets were snap-frozen in liquid nitrogen and stored at −20°C. Cells were lysed in 100 µl lysis buffer (5 ml contain: 10 mM Tris-HCl pH 7.5, 4% SDS, 1 PhosSTOP tablet [Roche], a scoop of DNase I [NEB]) for 5 min at room temperature. After spinning the samples for 5 min at max speed at room temperature in a table-top centrifuge, 80 µl of the cell lysate was taken up in 120 µl 1.6X SDS sample buffer (0.1 M Tris pH 6.8, 5% [v/v] glycerol, 0.2% [w/v] SDS, 1% [v/v] β-mercaptoethanol, 0.025% [w/v] bromophenol blue) and kept on ice. Subsequently, 20 µl of sample were run on 12% SDS-PAGE gels supplemented with 100 µM MnCl_2_ and 50 µM Phos-tag acrylamide compound (Wako) for 4 to 5 hours at 100 V at 4°C. Subsequent immunoblotting was performed as described below.

### Immunoblotting

Samples from mixed cultures grown in PYE were taken in exponential phase (OD_660_ of 0.3-0.4) and cells were harvested by centrifugation (max speed, 1 min, room temperature) in a table-top centrifuge. Sampling of synchronized cultures is described above. Pellets were resuspended in 1X SDS sample buffer (62.5 mM Tri-HCl pH 6.8, 10% [v/v] glycerol, 2% [w/v] SDS, 5% [v/v] β-mercaptoethanol, 0.001% [w/v] bromophenol blue) and boiled for 5-10 min before being run on 10-12.5% SDS-PAGE or precast Mini-Protean TGX (Biorad) gels. Proteins were transferred from SDS-PAGE gels to PVDF-membranes (Immobilon-P, Millipore 0.45µm) in transfer buffer (per 1 l: 3.03 g Tris base, 14.4 g glycine, 20% [v/v] ethanol, ddH2O) using a Biorad semi-dry system (24 V, 300 mA, 30 minutes) (Fig. 5A) or a Biorad wet blot system (100 V, 1 h, 4°C) (Fig. 7). For immunoblotting with Phos-tag gels, gels were kept at 4°C and washed successively for 10 min with transfer buffer containing 10 mM EDTA and transfer buffer containing 0.1% (w/v) SDS, and transfer to PVDF-membranes (Immobilon-P, Millipore, 0.45µm) was done using a Biorad wet blot system (80 V, 120 min, 4°C). After transfer, membranes were blocked by incubation with blocking buffer for 1 h at room temperature. Blocking buffer was 1X PBS (137 mM NaCl, 27 mM KCl, 81 mM Na_2_HPO_4_, 18 mM KH_2_PO_4_) containing 0.1% (v/v) Tween20 and 5% (w/v) skimmed milk powder (for experiments shown in Fig. 7) or TBST (20 mM 1 M Tris-HCl pH 7.5, 150 mM 5 M NaCl, 0.1% [v/v] Tween20) containing 5% skimmed milk powder (all other experiments). After blocking, membranes were incubated with the primary antibody in blocking buffer for 1 h at room temperature or overnight at 4°C, followed by three washes in blocking buffer (5 min, room temperature) and incubation with the secondary antibody in blocking buffer for 1 h at room temperature. Membranes were washed three to four times with blocking buffer and three times in 1X PBS (for experiments shown in Fig. 7) before addition of ECL detection reagent (KPL LumiGLO or LumiGLO Reserve, SeraCare Life Sciences). Chemiluminescence was detected using a Fujifilm LAS-4000 Imager (Fuji) with automatic exposure time determination or by exposure of membranes to X-ray films (Fujifilm). Primary antibodies were used at the following dilutions: α-MreB (1:20’000) (a gift from Régis Hallez), α-TacA (1:15’000) ^12^, α-SpmX (1:10’000) ^12^, α-PleC (1:5’000) ^31^, α-CtrA (1:5’000) ^32^, M2 α-FLAG (1:10’000) (Sigma). Secondary antibodies HRP-conjugated rabbit anti-mouse and swine anti-rabbit (Dako Cytomation, DK) were used at a 1:10’000 dilution.

### Microscopy and Image analysis

Cultures were grown in PYE and imaged in exponential phase (OD_660_ of 0.3-0.4) on 1% agarose PYE pads. Microscopy images were acquired using softWoRx 6.0 (GE Healthcare) on a DeltaVision system (GE Healthcare), equipped with a pco.edge sCMOS camera, and an UPlan FL N 100X/1.30 oil objective (Olympus). DivJ-mCherry localization was analyzed using Fiji software package ^33^ with the MicrobeJ plugin ^34^. Oufti ^35^ was used to quantify cell length and data were analyzed in Prism 7 (GraphPad) with statistical testing using ordinary one-way ANOVA and Tukey’s multiple comparision test. SpmX-mCherry localization and stalk formation (Supplementary Fig. 1) was analysed manually.

### Time-lapse microscopy and Image analysis

For Figs. 5 and Supplementary Fig. 6 *spmX* promoter activity during the cell cycle of single cells of *C. crescentus* wild type and selected mutants was analyzed using P*spmX*-Dendra2 and time-lapse microscopy. Strains were grown overnight in 5 ml PYE supplemented with appropriate antibiotics in a roller drum at 30°C. On the day of the experiment, these cultures were diluted 50-fold in 5 ml PYE supplemented with 2.5 µg/ml oxytetracycline and if appropriate with 0.1 mM IPTG and allowed to grow in a roller drum at 30°C until they reached an OD_660_ of 0.2. Cells were mounted on 1% PYE medium agar pads containing appropriate supplements sealed with a double layer of gene frames (Thermo Fisher Scientific; 1.7 x 2.8 cm) and subjected to time-lapse microscopy using softWoRx 6.0 (GE Healthcare) on a DeltaVision system (GE Healthcare), equipped with a pco.edge sCMOS camera, and an UPlan FL N 100X/1.30 oil objective (Olympus) using 0.3 s exposure time for phase-contrast, FITC and TRITC channels and a frame rate of 1 frame/15 minutes for different amounts of time. For Figs. 5a-c, Supplementary Fig. 6, green Dendra2 was irreversibly photoconverted to red in the latest detectable pre-divisional stage 15 minutes before the first visible cell separation and birth of the swarmer cell using 2 seconds UV light (DAPI channel) leading to a conversion efficiency from green to red between 62-75%. Image processing and analysis was done using the Fiji software package ^33^. From every pre-divisional cell and its offspring (swarmer and stalked cell) the total cell fluorescence and cell area was manually measured for every time-point over the course of one cell cycle and if appropriate for up to three consecutive cell cycles and the integrated density (mean intensity normalised to cell area) determined. In case of time-point 0 (predivisional cell after photoconversion), the mean intensity of the pre-divisional was normalised to the area of the swarmer or stalked cell. Integrated density values were then normalised for background fluorescence to obtain normalised fluorescence values over the cell cycle, plotted as arbitrary units (A.U.) in Fig. 5c.

In order to determine the fraction of cells with induced *spmX* promoter activity in Fig. 5c, for individual cells the mean of all normalised fluorescence values over one cell cycle was calculated. If the mean normalised fluorescence was above 150 A.U., cells were scored as showing induced P*spmX* activity; if below 150 A.U. then cells were scored as non-induced.

In Supplementary Fig. 6 normalised fluorescence values over the cell cycle of each cell type were set relative to the value of the late predivisional cell after photoconversion (t0=1). Green Dendra2 was irreversibly photoconverted to red only once in the latest detectable pre-divisional stage 15 minutes before the first visible cell separation and birth of the swarmer cell of generation 1. Image analysis was done as described above. Normalised fluorescence values of each cell type over each of the three cell cycles were set relative to the first late predivisional cell after photoconversion. The slope of relative fluorescence was determined over the course of one cell cycle and cells were scored as showing induced activity if the slope ≥ 0.19 or as not showing activity if the slope ≤ 0.19. The fraction of cells with induced *spmX* promoter activity was then calculated and plotted.

The ON kinetics of P*spmX* activity in Fig. 5b was determined as following: Dendra2 was photoconverted once using 2 seconds UV light (DAPI channel) in the latest detectable pre-divisional stage 15 minutes before the first visible cell separation and birth of the swarmer cell. Cells were normalized and evaluated as above. The time of the first visible increase of cell fluorescence in the swarmer cell (green line) and the stalked cell (grey line) cells with slope ≥ 0.075 was plotted as fraction of cells with induced *spmX* promoter activity over time. For inferred OFF kinetics (blue dashed line), cells at different cell cycle stages were photoconverted once using 2 seconds UV light (DAPI channel), normalized as above and the increase of fluorescence within 15-minute intervals was determined using the first time-point after visible cell separation as reference. Fraction of cells scored as having induced *spmX* promoter activity over the cell cycle was plotted over time. Results were plotted using Prism 7 (GraphPad).

### β-Galactosidase assays

For *β*-galactosidase assays strains harboring pAK502-based *lacZ* reporter plasmids were grown in 2 ml PYE supplemented with chloramphenicol and additional antibiotics as appropriate overnight at 30°C in a drum roller and diluted the next day 20-fold in 2 ml of the same medium, followed by further incubation for an additional 4.5 h under the same conditions before sampling. *β*-Galactosidase assays were performed as described before ^36^. For *β*-galactosidase assays strains harboring pRKlac290-based plasmids were grown overnight in 5 ml PYE supplemented with the appropriate antibiotics in a roller drum at 30°C. On the day of the experiment, these cultures were diluted 20-fold in 5 ml PYE supplemented with appropriate antibiotics and allowed to grow in a roller drum at 30°C until they reached an OD_660_ of 0.3 to 0.54. 1.8 ml culture was spun and the pellet resuspended in 1.8 ml Z-buffer (0.06 M Na2HPO4, 0.04 M NaH2PO4, 0.01 M KCl, 0.001 M MgSO_4_, 0.05 M β-mercaptoethanol) of which 0.8 ml were used for OD_660_ measurements and 1 ml transferred to a chloroform-stable Eppendorf tube. Subsequently, 100 µl 0.1% SDS and 20 µl chloroform were added. Samples were vortexed for 10 sec and incubated at room temperature for 30-60 min. From the top aqueous layer of the mixture three times 200 µl (three technical replicates) were transferred to a 96-well plate, 25 µl ONPG (4 mg/ml in Z-buffer) were added to each well and consumption of ONPG was followed at 405 nm using a BioTek Instruments EL800 plate reader (20 reads at the fastest interval). β-galactosidase activity was calculated as the initial slope of the increase of OD405/time at a linear range and corrected for the OD_660_ and volume. Activities were normalized to the activity of a wild-type strain that was included in all assays.

### Flow cytometry

Cultures were inoculated from a fresh colony and strains were grown in 2 ml PYE at 30°C in a drum roller 20 h (stationary phase) or diluted back 20-fold in 2 ml of the same medium after overnight incubation, followed by incubation for an additional 4.5 h (exponential phase). Rifampicin was added to cultures at a final concentration of 25 ug/ml and cultures were incubated for 2 h under the same conditions before cells were being fixed in cold 70% ethanol. Cells were collected by centrifugation, resuspended in 0.5 ml FACS buffer (10 mM Tris-HCl pH 7.5, 1 mM EDTA, 50 mM sodium citrate, 0.01% [v/v] TritonX-100) containing 2.5 µl RNaseA solution (Sigma) and incubated at room temperature for 30 min, after which 50 µl were transferred to 1 ml FACS buffer containing 1.5 µl YO-PRO-1 iodide (Thermo Fisher Scientific) and incubated for 2 h at room temperature in the dark. Data were acquired using a FACS Canto II (BD Biosciences) with 50’000 events recorded and analyzed with FlowJo software (FlowJo LLC).

### Mass spectrometry-based proteome analysis

10^9^ *C. crescentus* cells grown in PYE to exponential phase were collected, washed twice with phosphate buffer, resuspended in 50 µl lysis buffer (1% sodium deoxycholate, 0.1M ammoniumbicarbonate), reduced with 5 mM TCEP for 15 min at 95°C and alkylated with 10 mM chloroacetamide for 30 min at 37°C. Samples were digested with trypsin (Promega) at 37°C overnight (protein to trypsin ratio: 50:1) and desalted on C18 reversed phase spin columns according to the manufacturer’s instructions (Microspin, Harvard Apparatus).

1 µg of peptides of each sample were subjected to LC–MS analysis using a dual pressure LTQ-Orbitrap Elite mass spectrometer connected to an electrospray ion source (both Thermo Fisher Scientific). Peptide separation was carried out using an EASY nLC-1000 system (Thermo Fisher Scientific) equipped with a RP-HPLC column (75 µm × 30 cm) packed in-house with C18 resin (ReproSil-Pur C18–AQ, 1.9µm resin; Dr. Maisch GmbH, Ammerbuch-Entringen, Germany) using a linear gradient from 95% solvent A (0.15% formic acid, 2% acetonitrile) and 5% solvent B (98% acetonitrile, 0.15% formic acid) to 28% solvent B over 75min at a flow rate of 0.2 µl/min. The data acquisition mode was set to obtain one high resolution MS scan in the FT part of the mass spectrometer at a resolution of 240,000 full width at half-maximum (at m/z 400) followed by MS/MS scans in the linear ion trap of the 20 most intense ions. The charged state screening modus was enabled to exclude unassigned and singly charged ions and the dynamic exclusion duration was set to 20s. The ion accumulation time was set to 300 ms (MS) and 50 ms (MS/MS). The collision energy was set to 35%, and one microscan was acquired for each spectrum. For all LC-MS measurements, singly charged ions and ions with unassigned charge state were excluded from triggering MS2 events.

To determine changes in protein expressions across samples, a MS1-based label-free quantification was carried out. Therefore, the generated raw files were imported into the Progenesis QI software (Nonlinear Dynamics, Version 2.0) and analyzed using the default parameter settings. MS/MS-data were exported directly from Progenesis QI in mgf format and searched against a decoy database of the forward and reverse sequences of the predicted proteome from *Caulobacter crescentus* (strain NA1000 / CB15N) (Uniprot, download date: 08/09/2015, total of (8,234 entries) using MASCOT. The search criteria were set as following: full tryptic specificity was required (cleavage after lysine or arginine residues); 3 missed cleavages were allowed; carbamidomethylation (C) was set as fixed modification; oxidation (M) as variable modification. The mass tolerance was set to 10 ppm for precursor ions and to 0.6 Da for fragment ions. Results from the database search were imported into Progenesis QI and the final peptide measurement list containing the peak areas of all identified peptides, respectively, was exported. This list was further processed and statically analyzed using our in-house developed SafeQuant R script (SafeQuant, https://github.com/eahrne/SafeQuant, ^37^). The peptide and protein false discovery rate (FDR) was set to 1% using the number of reverse hits in the dataset. All quantitative analyses were performed in biological triplicates. All raw data and results associated with the manuscript have been deposited to the ProteomeXchange Consortium via the PRIDE ^38^ partner repository with the dataset identifier PDX012739 (Reviewer account details: username, password: F3it2yqf).

### Genetic selection for c-di-GMP-independent mutations in *shkA*

Six independent colonies of strain AKS1 (CB15 rcdG^0^ Δ*lacA*::Ω *ampG*::pNPTStet-ampG) harboring in addition plasmid pAK503-spmX were inoculated into 5 ml PYE containing chloramphenicol (1 mg/l) and oxytetracycline (10 mg/l) and grown overnight. For UV mutagenesis, 2 ml of each culture was distributed in 6-well plates and irradiated with 10’000 mJ UV light using a Stratalinker, after which 5 ml PYE containing chloramphenicol (1 mg/l) and oxytetracycline (10 mg/l) were added and cultures grown for 7h at 30°C with shaking. 500µl of each culture were plated on PYE plates containing chloramphenicol (1 mg/l), oxytetracycline (10 mg/l) and kanamycin (6.25 mg/l) and plates incubated at 30°C for 4 days. All colonies from one plate were pooled, diluted to an OD_660_ of 0.1 and grown for 7h at 30°C, and phage lysates were prepared for each independent pool. Following transduction of strain AKS17 (CB15 rcdG^0^ Δ*lacA*::Ω /pAK502-spmX) with individual pool lysates, cells were plated on PYE containing chloramphenicol (1 mg/l), oxytetracycline (10 mg/l) and X-Gal (40 mg/l) and incubated at 30°C for 3 days. Blue colonies were once re-streaked to confirm their blue colony phenotype, followed by colony PCR using primers 9765 and 9766 and sequencing (Microsynth, Balgach, Switzerland) of the PCR product using primers 9765, 9766, 6011 and 6498.

### Genetic screen for mutations abolishing *spmX’-‘lacZ* activity

Transposon mutagenesis was performed by delivery of pSAM-Ec in strain UJ6168 carrying pAK502-spmX by conjugation and cells were plated on PYE containing chloramphenicol (1 mg/l), kanamycin (50 mg/l), nalidixic acid (20 mg/l) and X-Gal (40 mg/l) and incubated at 30°C for 3 days. White colonies were pooled in 5 ml PYE supplemented with appropriate antibiotics and grown overnight to prepare φCR30 lysates. From this pool lysate transposon insertions were transduced in a clean background (UJ6168/pAK502-spmX), followed by blue/white screening as described above. Lysates were prepared from single white colonies and mutations transduced in a NA1000 wild-type background. Transposon insertion sites were mapped using a two-step semi-arbitrary PCR approach as described before ^39^. Primers 9058, 9003 and 9004 were used in the first PCR and primers 9058 and 9005 in the second PCR. PCR products were sequenced (Microsynth, Balgach, Switzerland) with primer 9058. In one strain (AKS50), the transposon mapped within *pleD* at position 2’695’825 of the NA1000 reference genome ^40^.

### Protein expression and purification

For isothermal titration calorimetry (ITC) and *in vitro* phosphorylation experiments shown in Fig. 2, proteins were expressed and purified as follows. *E. coli* Rosetta 2(DE3) cells were used to express proteins from pET28a and pET32b expression plasmids. Cells were grown in LB-Miller supplemented with the appropriate antibiotics to an OD600 of 0.4 to 0.6, expression was then induced with 0.5 mM IPTG for 4 h at 30°C. Proteins were purified on an ÄKTApurifier 10 system (GE Healthcare) using 1 ml HisTrap HP columns (GE Healthcare) followed by size exclusion chromatography (HiLoad 16/60 Superdex 200) using the following buffers: lysis buffer (wash buffer supplemented with protease inhibitor, DNase I [NEB]), wash buffer (20mM HEPES-KOH pH 8.0, 0.5 M NaCl, 10% glycerol, 20 mM imidazole, 1 mM DTT), elution buffer (20 mM HEPES-KOH pH 8.0, 0.5 M NaCl, 10% glycerol, 500 mM imidazole, 1 mM DTT), storage buffer (10 mM HEPES-KOH pH 8.0, 50 mM KCl, 10% glycerol, 0.1 mM EDTA, 1 mM DTT). MgCl_2_ was added to reaction mixtures immediately prior to experiments to a final concentration of 5 mM. For all other experiments using purified ShkA, mutant variants thereof and ShkA_REC1_, except NMR experiments (see below), proteins were expressed in *E. coli* BL21(DE3) grown in LB-Miller at 37°C in 500-ml cultures, with IPTG induction (1 mM) at an OD600 of 0.5 to 0.8 followed by incubation for 4 h. Cells were harvested by centrifugation (5000 g, 20 min, 4°C), washed once with 20 ml of 1X PBS, flash-frozen in liquid N_2_, and stored at −80°C until purification. For purification, the pellet was resuspended in 8 ml of buffer A (2X PBS containing 500 mM NaCl, 10 mM imidazole and 2 mM β-mercaptoethanol) supplemented with DNaseI (AppliChem) and Complete Protease inhibitor (Roche). After one passage through a French press cell, the suspension was ultra-centrifuged (100’000 g, 30 min, 4°C) and the supernatant was mixed with 800 µl of Ni-NTA slurry, pre-washed with buffer A, and incubated for 1-2 h on a rotary wheel at 4°C. Ni-NTA agarose was loaded on a polypropylene column and washed with at least 50 ml of buffer A, after which the protein was eluted with 2.5 ml of buffer A containing 500 mM imidazole. The eluate was immediately loaded on a PD-10 column pre-equilibrated with kinase buffer (10 mM HEPES-KOH pH 8.0, 50 mM KCl, 10% glycerol, 0.1 mM EDTA, 5 mM MgCl_2_, 5 mM β-mercaptoethanol). The protein was then eluted with 3.5 ml of kinase buffer and stored at 4°C until further use, usually no longer than one week. Uniformly [^13^C, ^15^N]-labeled protein for NMR studies was prepared by growing BL21(DE3) cells harboring pET28a-shkAREC1 in 1 l M9 minimal medium, with 1 g ^15^NH4Cl and 2 g [*U*-^13^C] glucose per liter medium, at 37°C. Expression was induced at an OD600 of 0.8 with 1 mM IPTG and cells were harvested 4 h post-induction. ShkA_REC1_ was purified using Ni-NTA slurry as described above, followed by size exclusion chromatography (HiLoad 16/60 Superdex 200) on a ÄKTApurifier 10 system (GE Healthcare) with NMR buffer (25 mM Tris pH 7.2 with 50 mM KCl, 2 mM MgSO_4_) and concentration of the sample using Vivaspin6 5 kDa MWCO concentrators (Sartorius).

### Isothermal titration calorimetry (ITC)

ITC binding assays were performed with a VP-ITC isothermal titration calorimeter (MicroCal). Concentrations were 12 µM ShkA in the cell and 130 µM cdG in the syringe. ITC Buffer: 10 mM HEPES-KOH pH 8.0, 100 mM NaCl, 10% glycerol, 1 mM DTT. After a first injection of 3 µl, 10 µl was injected at 29 time points. Data analysis was performed with MicroCal (ORIGIN) and fitted with the One binding site model of ORIGIN.

### *In vitro* phosphorylation assays

Kinase assays were adapted from ^9^. Reactions were performed in kinase buffer supplemented with 433 µM ATP and 5-20 µCi [γ^32^P]-ATP (3000Ci/mmol, Hartmann Analytic) at room temperature if not otherwise mentioned. Additional nucleotides were added as indicated in the Fig.s or Fig. legends. Kinase reactions were run for 3.5 min if not otherwise mentioned and stopped by addition of 5X SDS sample buffer and stored on ice, then run on 12.5% SDS-PAGE or precast Mini-Protean TGX (Biorad) gels. Wet gels were exposed to a phosphor screen for 0.5 h to 3 h and then scanned using a Typhoon FLA 7000 imaging system (GE Healthcare), after which gels were stained with Coomassie Brilliant Blue.

### C-di-GMP UV crosslinking assays

^32^P-labelled c-di-GMP was prepared by incubation of 250 µCi [α^32^P]-GTP (3000Ci/mmol, Hartmann Analytic) in 200 ul DgcZ reaction buffer (50 mM Tris-HCl pH 7.5, 50 mM NaCl, 5 mM MgCl_2_) with 2 µM purified DgcZ for 24 h at 30°C. DgcZ (YdeH) was purified as described previously ^41^. The reaction was stopped by boiling and, after centrifugation (1 min, 16’000 g), the supernatant containing radiolabeled c-di-GMP was removed and stored at −20°C. Non-radio-labelled c-di-GMP was prepared as described previously ^41^ with the following modifications. 5% (v/v) methanol and 5 mM tetraethylammonium bromide were added to the c-di-GMP reaction and the mixture was loaded on a Luna 10 µM C18(2) 100Å, 100 x 21.2 mm column (Phenomenex). GTP was removed by isocratic elution in TEAB buffer (5 mM tetraethylammonium bromide, 5% [v/v] methanol) over 3 column volumes (CV), after which c-di-GMP was eluted with a linear gradient from 0-20% ethanol in TEAB buffer over 4 CV. c-di-GMP was lyophilized overnight, resuspended in ddH2O at a concentration of 10 mM and aliquoted. Aliquots were lyophilized and stored at −20°C. ddH2O was added to aliquots to give a 10 mM working stock, the concentration of which was verified by measuring absorbance at 253 nm. A 80 mM stock solutions of hot c-di-GMP was prepared by mixing 10-20 µCi ^32^P-labeled c-di-GMP with 4 µl 1mM cold c-di-GMP in a final volume of 50 µl. More dilute stock solutions were prepared by mixing with ddH2O. UV crosslinking reactions (16 µl) contained 4 µl of hot c-di-GMP to give the indicated final concentrations and 0.5 µM ShkA, ShkA mutant variants or ShkA_REC1_ in kinase buffer, and were incubated at room temperature for 45 min before UV crosslinking at 4°C, 254 nm, 2 min on a Caprotec Protein Detector. For competition experiments, reactions contained a 50-fold molar excess of cold c-di-GMP over hot c-di-GMP added from a 10 mM stock. Samples were mixed with 5X SDS sample buffer, boiled for 10 min and run on 12.5% SDS-PAGE or precast Mini-Protean TGX (Biorad) gels. Gels were stained with Coomassie Brilliant Blue, dried and exposed to a phosphor screen that was scanned using a Typhoon FLA 7000 imaging system (GE Healthcare). Bands of autoradiographs were quantified using Fiji and normalized to wild-type ShkA, which was run in parallel to mutant variants or ShkA_REC1_ on the same gel for each experiment. Experiments were repeated twice and values represent means and standard deviations. Binding curves were fitted to a “Binding – saturation, One site – Total” model using Prism 7 (GraphPad).

### Nuclear magnetic resonance (NMR) spectroscopy

All NMR spectra were recorded at 25 °C on a Bruker Avance-700 spectrometer equipped with a cryogenically cooled triple-resonance probe. ShkA_REC1_ protein samples were prepared in 25 mM Tris pH 7.2 with 50 mM KCl and 2 mM MgSO_4_ in 95%/5% H2O/D2O. For the sequence-specific backbone resonance assignment of 950 μM [*U*-^15^N, ^13^C]-ShkA_REC1_, the following NMR experiments were recorded: 2D [^15^N,^1^H]-TROSY, 3D HNCA and 3D HNCACB. For the c-di-GMP binding experiments a series of 2D [^15^N,^1^H]-TROSY spectra of 200 μM [*U*-^15^N]-ShkA_REC1_ were recorded with c-di-GMP concentrations of 0 μM, 20 μM, 50 μM, 100 μM, 150 μM, 200 μM and 400 μM. Chemical shift perturbations (*Δδ*(HN)) of amide moieties were calculated as: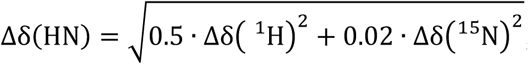, where Δδ.(^1^H) and Δδ.(^15^N), are the amide proton and amide nitrogen chemical shift differences to the reference spectrum, respectively. Combined secondary chemical shifts of C*α* and C*β* were calculated relative to the random-coil values of Kjaergaard and Poulson ^42^.

### C-di-GMP extraction and quantifications

Strains AKS371 and AKS412 were grown in 2 ml PYE overnight, diluted 40-fold the next day in 20 ml PYE and grown for 6h until reaching an OD_660_ of approximately 0.35. 14 ml were spun down (11’000 g, 5 min, 4°C), the pellet was washed once in 1 ml ddH2O (16’000 g, 2 min, 4°C) and snap-frozen in liquid nitrogen. Metabolites were extracted by resuspending the pellet in 300 ul ice-cold extraction solvent (40% [v/v] acetonitrile, 40% [v/v] methanol, 20% [v/v] ddH2O) and incubation at 95°C for 10 min. Extractions were repeated on the pellet twice with 200 ul extraction solvent and the pooled extracts (700 ul) were stored overnight at −80°C. Remaining debris was removed by centrifugation (16’000 g, 10 min, 4°C), the supernatant was transferred to a fresh microcentrifuge tube and solvent was removed using a Speedvac. LC-MS-based c-di-GMP quantification was performed as described previously ^43,44^ and cellular concentrations were calculated as described before ^11^. Experiments were performed in biological triplicates and values given are means and standard deviations.

### Sequence alignments, secondary structure prediction, structure modelling

The *C. crescentus* ShkA sequence was blasted against the non-redundant protein sequences (nr) database using default settings and the first 500 hits were aligned in Geneious using the Geneious Alignment algorithm with default settings (Supplementary Data 1). The alignment was manually checked for the presence of the DDR motif in the REC1-REC2 linker and the sequences that harbored this motif (see Fig. S3A) were realigned as described above. This alignment was used to create sequence logos using WebLogo3 ^27^, construct a phylogenetic tree using Geneious Tree Builder with default settings and calculate conservation scores using the ConSurf server ^45^. Four proteins of the original blast set that do not contain the DDR motif were included in the phylogenetic tree for comparison. Secondary structure prediction for *E. coli* BarA_PRD_ (residues 532-655) was performed with HHPred ^46^ and the ShkA_REC1_-BarA_PDR_ alignment was generated in Geneious as described above. The ShkAREC1-3GRC alignment was generated using HHPred. The ShkAREC1 structure was modelled using the SWISS-MODEL workspace ^47^ with PDB entry 3GRC as the template. Chemical shift perturbations and conservation scores mapped onto the ShkAREC1 model were visualized using MacPyMol (PyMOL v1.7.6.6, Schrödinger LLC).

## References for Supplementary Table S1

Abel, S. *et al.* (2013) ^11^

Radhakrishnan *et al.* (2008) ^12^

Aldridge *et al.* (2003) ^14^

Lori *et al.* (2018) ^36^

Alley *et al.* (1991) ^48^

Landgraf *et al.* (2012) ^49^

Thanbichler *et al.* (2007) ^50^

Wiles *et al.* (2013) ^51^

Roberts *et al.* (1996) ^52^

Matroule *et al.* (2004) ^53^

Kaczmarczyk *et al.* (2013) ^54^

Kaczmarczyk *et al.* (2012) ^55^

Collier *et al.* (2009) ^56^

## References for Supplementary Table S3

Abel *et al.* (2013) ^11^

Radhakrishnan *et al.* (2008) ^12^

Aldridge *et al.* (2003) ^14^

Alley *et al.* (1991) ^48^

Grant *et al.* (1990) ^57^

Vargas *et al.* (1999) ^58^

Woodcock *et al.* (1989) ^59^

McGrath *et al.* (2006) ^60^

Skerker *et al.* (2005) ^61^

Arellano *et al.* (2010) ^62^

Sprecher *et al.* (2017) ^63^

Agabian-Keshishian and Shapiro (1970) ^64^

Ely and Johnson (1977) ^65^

