## Supplementary Methods for "Precise transcription timing by a second-messenger drives a bacterial G1/S cell cycle transition"

**Plasmid construction details**

pAK503: *nptII* without RBS and start codon was PCR-amplified from pAK405 using primers 8959/9495, the product was digested with *Bsr*GI/*Xba*I and cloned into pAK501 cut with *Acc*65I/*Xba*I.

pAK502-*tacA*: The *spmX* promoter and part of the codon sequence were PCR-amplified from *C. crescentus* gDNA using primers 10015/10016, the product was digested with *Xba*I/*Kpn*I and cloned into pAK502 digested with the same enzymes.

pAK502-*spmX*: The *spmX* promoter and part of the codon sequence were PCR-amplified from *C. crescentus* gDNA using primers 8966/9064, the product was digested with *Xba*I/*Kpn*I and cloned into pAK502 digested with the same enzymes.

pAK503-*spmX*: The *spmX* promoter and part of the codon sequence were PCR-amplified from *C. crescentus* gDNA using primers 8966/9064, the product was digested with *Xba*I/*Kpn*I and cloned into pAK503 digested with the same enzymes.

pNPTStet: *tetA* and *tetR* were PCR-amplified from pQF using primers 9552/9553, the product was digested with *Spe*I/*Nde*I and cloned into pNPTS138 digested with *Ase*I/*Xba*I.

pNPTStet-*ampG*: Part of *ampG* located downstream of *shkA* was PCR-amplified from *C. crescentus* gDNA with primers 9554/9555, the product was digested with *Spe*I/*Kpn*I and cloned into pNPTStet digested with the same enzymes.

pET28a-*shkA*: *shkA* was PCR-amplified from *C. crescentus* gDNA with primers 9745/9746, the product was digested with *Nde*I/*Eco*RI and cloned into pET28a digested with the same enzymes.

pET28a-*shkA(D369N)*: *shkA(D369N)* was amplified from *C. crescentus* strain UJ9618 by colony-PCR using primers 9745/9746, the product was digested with *Nde*I/*Eco*RI and cloned into pET28a digested with the same enzymes.

pQF-*shkA*: *shkA* was PCR-amplified from *C. crescentus* gDNA with primers 9745/9746, the product was digested with *Nde*I/*Eco*RI and cloned into pQF digested with *Ase*I/*Eco*RI.

pQF-*shkA(D369N)*: *shkA(D369N)* was amplified from *C. crescentus* strain UJ9618 by colony-PCR using primers 9745/9746, the product was digested with *Nde*I/*Eco*RI and cloned into pQF digested with *Ase*I/*Eco*RI.

pQF-*shkA(R26A)*: The mutant allele was generated by SOE-PCR using pET28a-*shkA* as template and flanking primers 669/670 and mutagenic primers 9904/9905. The product was digested with *Nde*I/*Eco*RI and cloned into pQF digested with *Ase*I/*Eco*RI.

pQF-*shkA(H61A)*: The mutant allele was generated by SOE-PCR using pET28a-*shkA* as template and flanking primers 669/670 and mutagenic primers 9968/9969. The product was digested with *Nde*I/*Eco*RI and cloned into pQF digested with *Ase*I/*Eco*RI.

pQF-*shkA(R74A)*: The mutant allele was generated by SOE-PCR using pET28a-*shkA* as template and flanking primers 669/670 and mutagenic primers 9970/9971. The product was digested with *Nde*I/*Eco*RI and cloned into pQF digested with *Ase*I/*Eco*RI. T*His shkA* allele contains a second mutation, A451V.

pQF-*shkA(W113A)*: The mutant allele was generated by SOE-PCR using pET28a-*shkA* as template and flanking primers 669/670 and mutagenic primers 9906/9907. The product was digested with *Nde*I/*Eco*RI and cloned into pQF digested with *Ase*I/*Eco*RI. T*His shkA* allele contains a second mutation, A451V.

pQF-*shkA(D125N)*: The mutant allele was generated by SOE-PCR using pET28a-*shkA* as template and flanking primers 669/670 and mutagenic primers 10009/10010. The product was digested with *Nde*I/*Eco*RI and cloned into pQF digested with *Ase*I/*Eco*RI.

pQF-*shkA(R128A)*: The mutant allele was generated by SOE-PCR using pET28a-*shkA* as template and flanking primers 669/670 and mutagenic primers 10005/10006. The product was digested with *Nde*I/*Eco*RI and cloned into pQF digested with *Ase*I/*Eco*RI.

pQF-*shkA(R130A)*: The mutant allele was generated by SOE-PCR using pET28a-*shkA* as template and flanking primers 669/670 and mutagenic primers 10007/10008. The product was digested with *Nde*I/*Eco*RI and cloned into pQF digested with *Ase*I/*Eco*RI.

pQF-*shkA(R128A, R130A)*: The mutant allele was generated by SOE-PCR using pET28a-*shkA* as template and flanking primers 669/670 and mutagenic primers 9908/9909. The product was digested with *Nde*I/*Eco*RI and cloned into pQF digested with *Ase*I/*Eco*RI.

pQF-*shkA(R177A)*: The mutant allele was generated by SOE-PCR using pET28a-*shkA* as template and flanking primers 669/670 and mutagenic primers 9910/9911. The product was digested with *Nde*I/*Eco*RI and cloned into pQF digested with *Ase*I/*Eco*RI. T*His shkA* allele contains a s*Eco*nd mutation, P233L.

pQF-*shkA(F230A, F234A)*: The mutant allele was generated by SOE-PCR using pET28a-*shkA* as template and flanking primers 669/670 and mutagenic primers 9972/9973. The product was digested with *Nde*I/*Eco*RI and cloned into pQF digested with *Ase*I/*Eco*RI.

pQF-*shkA(D297N)*: The mutant allele was generated by SOE-PCR using pET28a-*shkA* as template and flanking primers 669/670 and mutagenic primers 9974/9975. The product was digested with *Nde*I/*Eco*RI and cloned into pQF digested with *Ase*I/*Eco*RI.

pQF-*shkA(P321A)*: The mutant allele was generated by SOE-PCR using pET28a-*shkA* as template and flanking primers 669/670 and mutagenic primers 10013/10014. The product was digested with *Nde*I/*Eco*RI and cloned into pQF digested with *Ase*I/*Eco*RI.

pQF-*shkA(R324A)*: The mutant allele was generated by SOE-PCR using pET28a-*shkA* as template and flanking primers 669/670 and mutagenic primers 9912/9913. The product was digested with *Nde*I/*Eco*RI and cloned into pQF digested with *Ase*I/*Eco*RI.

pQF-*shkA(D325N)*: The mutant allele was generated by SOE-PCR using pET28a-*shkA* as template and flanking primers 669/670 and mutagenic primers 10011/10012. The product was digested with *Nde*I/*Eco*RI and cloned into pQF digested with *Ase*I/*Eco*RI.

pQF-*shkA(I327A)*: The mutant allele was generated by SOE-PCR using pET28a-*shkA* as template and flanking primers 669/670 and mutagenic primers 9976/9977. The product was digested with *Nde*I/*Eco*RI and cloned into pQF digested with *Ase*I/*Eco*RI. T*His shkA* allele contains a second mutation, A451V.

pQF-*shkA(Y338A)*: The mutant allele was generated by SOE-PCR using pET28a-*shkA* as template and flanking primers 669/670 and mutagenic primers 9978/9979. The product was digested with *Nde*I/*Eco*RI and cloned into pQF digested with *Ase*I/*Eco*RI.

pQF-*shkA(R344A)*: The mutant allele was generated by SOE-PCR using pET28a-*shkA* as template and flanking primers 669/670 and mutagenic primers 9914/9915. The product was digested with *Nde*I/*Eco*RI and cloned into pQF digested with *Ase*I/*Eco*RI. T*His shkA* allele contains a second mutation, A451V.

pQF-*shkA(E402A, G403A)*: The mutant allele was generated by SOE-PCR using pET28a-*shkA* as template and flanking primers 669/670 and mutagenic primers 9980/9981. The product was digested with *Nde*I/*Eco*RI and cloned into pQF digested with *Ase*I/*Eco*RI.

pQF-*shkA(C404A)*: The mutant allele was generated by SOE-PCR using pET28a-*shkA* as template and flanking primers 669/670 and mutagenic primers 9986/9987. The product was digested with *Nde*I/*Eco*RI and cloned into pQF digested with *Ase*I/*Eco*RI.

pQF-*shkA(D430N)*: The mutant allele was generated by SOE-PCR using pET28a-*shkA* as template and flanking primers 669/670 and mutagenic primers 10028/10029. The product was digested with *Nde*I/*Eco*RI and cloned into pQF digested with *Ase*I/*Eco*RI.

pQF-*shkA(D465A)*: The mutant allele was generated by SOE-PCR using pET28a-*shkA* as template and flanking primers 669/670 and mutagenic primers 9982/9983. The product was digested with *Nde*I/*Eco*RI and cloned into pQF digested with *Ase*I/*Eco*RI. This *shkA* allele contains a second mutation, *A451V*.

pQF-*shkA(C469A)*: The mutant allele was generated by SOE-PCR using pET28a-*shkA* as template and flanking primers 669/670 and mutagenic primers 9984/9985. The product was digested with *Nde*I/*Eco*RI and cloned into pQF digested with *Ase*I/*Eco*RI. This *shkA* allele contains a second mutation, *A451V*.

pQF-*shkA(K341)*: The mutant allele was generated by SOE-PCR using pET28a-*shkA* as template and flanking primers 669/670 and mutagenic primers 11171/11172. The product was digested with *Nde*I/*Eco*RI and cloned into pQF digested with *Ase*I/*Eco*RI.

pQF-*shkA(P342A)*: The mutant allele was generated by SOE-PCR using pET28a-*shkA* as template and flanking primers 669/670 and mutagenic primers 11173/11174. The product was digested with *Nde*I/*Eco*RI and cloned into pQF digested with *Ase*I/*Eco*RI.

pQF-*shkA(L343A)*: The mutant allele was generated by SOE-PCR using pET28a-*shkA* as template and flanking primers 669/670 and mutagenic primers 11175/11176. The product was digested with *Nde*I/*Eco*RI and cloned into pQF digested with *Ase*I/*Eco*RI.

pQF-*shkA(I340A)*: The mutant allele was generated by SOE-PCR using pET28a-*shkA* as template and flanking primers 669/670 and mutagenic primers 11177/11178. The product was digested with *Nde*I/*Eco*RI and cloned into pQF digested with *Ase*I/*Eco*RI.

pQF-*shkA(L338A)*: The mutant allele was generated by SOE-PCR using pET28a-*shkA* as template and flanking primers 669/670 and mutagenic primers 11179/11180. The product was digested with *Nde*I/*Eco*RI and cloned into pQF digested with *Ase*I/*Eco*RI.

pQF-*shkA(G337A)*: The mutant allele was generated by SOE-PCR using pET28a-*shkA* as template and flanking primers 669/670 and mutagenic primers 11181/11182. The product was digested with *Nde*I/*Eco*RI and cloned into pQF digested with *Ase*I/*Eco*RI.

pQF-*shkA(K331A)*: The mutant allele was generated by SOE-PCR using pET28a-*shkA* as template and flanking primers 669/670 and mutagenic primers 11183/11184. The product was digested with *Nde*I/*Eco*RI and cloned into pQF digested with *Ase*I/*Eco*RI.

pQF-*shkA(S347A)*: The mutant allele was generated by SOE-PCR using pET28a-*shkA* as template and flanking primers 669/670 and mutagenic primers 11185/11186. The product was digested with *Nde*I/*Eco*RI and cloned into pQF digested with *Ase*I/*Eco*RI.

pQF-*shkA(Q351A)*: The mutant allele was generated by SOE-PCR using pET28a-*shkA* as template and flanking primers 669/670 and mutagenic primers 11187/11188. The product was digested with *Nde*I/*Eco*RI and cloned into pQF digested with *Ase*I/*Eco*RI.

pQF-*shkA(D369N, H61A)*: A fragment harboring part of *shkA* was released from pQF-*shkA(H61A)* by digestion with *Spe*I/*Pst*I and cloned into pQF-*shkA(D369N)* digested with the same enzymes.

pQF-*shkA(D369N, R128A)*: A fragment harboring part of *shkA* was released from pQF-*shkA(R128A)* by digestion with *Spe*I/*Pst*I and cloned into pQF-*shkA(D369N)* digested with the same enzymes.

pQF-*shkA(D369N, R130)*: A fragment harboring part of *shkA* was released from pQF-*shkA(R130A)* by digestion with *Spe*I/*Pst*I and cloned into pQF-*shkA(D369N)* digested with the same enzymes.

pQF-*shkA(D369N, R128A, R130A)*: A fragment harboring part of *shkA* was released from pQF-*shkA(R128A, R130A)* by digestion with *Spe*I/*Pst*I and cloned into pQF-*shkA(D369N)* digested with the same enzymes.

pQF-*shkA(D369N, F230A, F234A)*: A fragment harboring part of *shkA* was released from pQF-*shkA(F230A, F234A)* by digestion with *Spe*I/*Pst*I and cloned into pQF-*shkA(D369N)* digested with the same enzymes.

pQF-*shkA(D369N, R128A)*: A fragment harboring part of *shkA* was released from pQF-*shkA(R128A)* by digestion with *Spe*I/*Pst*I and cloned into pQF-*shkA(D369N)* digested with the same enzymes.

pQF-*shkA(D369N, R324A)*: A fragment harboring part of *shkA* was released from pQF-*shkA(R324A*) by digestion with *Spe*I/*Pst*I and cloned into pQF-*shkA(D369N)* digested with the same enzymes.

pQF-*shkA(D369N, Y338A)*: A fragment harboring part of *shkA* was released from pQF-*shkA(Y338A)* by digestion with *Spe*I/*Pst*I and cloned into pQF-*shkA(D369N)* digested with the same enzymes.

pET28a-*shkA(D430N)*: The mutant allele was generated by SOE-PCR using pET28a-*shkA* as template and flanking primers 669/670 and mutagenic primers 10028/10029. The product was digested with *Nde*I/*Eco*RI and cloned into pET28a digested with *Nde*I/*Eco*RI.

pET28a-*shkA(E386Q, D387N)*: The mutant allele was generated by SOE-PCR using pET28a-*shkA* as template and flanking primers 669/670 and mutagenic primers 10026/10027. The product was digested with *Nde*I/*Eco*RI and cloned into pET28a digested with *Nde*I/*Eco*RI.

pET28a-*shkA(K480R)*: The mutant allele was generated by SOE-PCR using pET28a-*shkA* as template and flanking primers 669/670 and mutagenic primers 10024/10025. The product was digested with *Nde*I/*Eco*RI and cloned into pET28a digested with *Nde*I/*Eco*RI.

pET28a-*shkA(R324A)*: A fragment harboring part of *shkA* was released from pQF-*shkA(R324A)* by digestion with *Asc*I/*Pst*I and cloned into pET28a-*shkA* digested with the same enzymes.

pET28a-*shkA(D369N, R324A)*: A fragment harboring part of *shkA* was released from pQF-*shkA(R324A)* by digestion with *Asc*I/*Pst*I and cloned into pET28a-*shkA(D369N)* digested with the same enzymes.

pET28a-*shkA(Y338A)*: A fragment harboring part of *shkA* was released from pQF-*shkA(Y338A)* by digestion with *Asc*I/*Pst*I and cloned into pET28a-*shkA* digested with the same enzymes.

pET28a-*shkA(D369N, Y338A)*: A fragment harboring part of *shkA* was released from pQF-*shkA(Y338A)* by digestion with *Asc*I/*Pst*I and cloned into pET28a-*shkA(D369N)* digested with the same enzymes.

pET28a-*shkA(R128A, R130A)*: The mutant allele was generated by SOE-PCR using pET28a-*shkA* as template and flanking primers 669/670 and mutagenic primers 9908/9909. The product was digested with *Nde*I/*Eco*RI and cloned into pET28a digested with the same enzymes.

pET28a-*shkA-REC1*: A fragment of *shkA* encoding REC1 was PCR-amplified from *C. crescentus* gDNA with primers 10862/10866, the product was digested with *Nde*I/*Kpn*I and cloned into pUC18 digested with the same enzymes. A *Nde*I/*Eco*RI from the resulting plasmid was subcloned into pET28a digested with the same enzymes.

pQF-*pleC*: A DNA fragment including *pleC* ORF and the mapped *pleC* promoter as PCR-amplified from plasmid pJS14-*pleC* with primers 10781/10783, the product was digested with *Spe*I/*Kpn*I and cloned into pQF digested with the same enzymes.

pQF-*pleC(K-P-)*: A DNA fragment including *pleC(T614R)* ORF and the mapped *pleC* promoter as PCR-amplified from plasmid pJS14-*pleC(T615R)* with primers 10781/10783, the product was digested with *Spe*I/*Kpn*I and cloned into pQF digested with the same enzymes.

pQF-*pleC(K-P+)*: A DNA fragment including *pleC*(F778L) ORF and the mapped *pleC* promoter as PCR-amplified from plasmid pJS14-*pleC*(F778L) with primers 10781/10783, the product was digested with *Spe*I/*Kpn*I and cloned into pQF digested with the same enzymes.

p*divJ-mCherry*: The 3' part of *divJ* was PCR-amplified from *C. crescentus* gDNA with primers 10205/10206, the product was digested with *Kpn*I/*Age*I and cloned into pCHYC-4 digested with the same enzymes.

pNPTS138-Δ*tacA*: Two DNA fragments flanking the *tacA* ORF were PCR-amplified from *C. crescentus* gDNA. Fragment 1 was amplified with primers 6535/6098, then digested with *Pst*I/*Kpn*I. Fragment 2 was amplified with primers 6099/6536, then digested with *Kpn*I/*Eco*RI. The two fragments were ligated to pNPTS138 digested with *Pst*I/*Eco*RI.

pNPTS138-Δ*shkA*: Two DNA fragments flanking the *shkA* ORF were PCR-amplified from *C. crescentus* gDNA. Fragment 1 was amplified with primers 6085/6086, then digested with *Pst*I/*Kpn*I. Fragment 2 was amplified with primers 6087/6088, then digested with *Kpn*I/*Eco*RI. The two fragments were ligated to pNPTS138 digested with *Pst*I/*Eco*RI.

pNPTS138-*3xFLAG-tacA*: The upstream region of the *tacA* ORF was PCR-amplified from *C. crescentus* gDNA with primers 6438/6439 (fragment 1). A 5' part of *tacA* was PCR-amplified from *C. crescentus* gDNA with primers 6440/7604 (fragment 2), and then used as a template with primers 6441/7604 to insert the coding sequence of the 3xFlag-Tag (fragment 3). SOE-PCR was performed with fragment 1 and 3 as template and primers 6438/7604. The resulting product was then digested with *Pst*I/*Eco*RI and ligated into pNPTS138 digested with the same enzymes.

pNPTS138-*3xFLAG shkA*: The upstream region of the *shkA* ORF was PCR-amplified from *C. crescentus* gDNA with primers 6442/6443 (fragment 1). A 5' part of *shkA* was PCR-amplified from *C. crescentus* gDNA with primers 6444/7605 (fragment 2), and then used as a template with primers 6445/7605 to insert the coding sequence of the 3xFlag-Tag (fragment 3). SOE-PCR was performed with fragment 1 and 3 as template and primers 6442/7605. The resulting product was then digested with *Pst*I/*Eco*RI and ligated into pNPTS138 digested with the same enzymes.

pNPTS138-*shkA(DD)*: The 5' region of *shkA* was PCR-amplified from *C. crescentus* gDNA with primers 8169/8170 (fragment 1). The downstream region of the *shkA* ORF was PCR-amplified from *C. crescentus* gDNA with primers 8171/8172 (fragment 2). SOE-PCR was performed with the two fragments as template and primers 8169/8172. The resulting product was then digested with *Bam*HI/*Eco*RI and ligated into pNPTS138 digested with the same enzymes.

pNPTS138-*tacA(DD)*: The 5' region of *tacA* was PCR-amplified from *C. crescentus* gDNA with primers 7925/7716 (fragment 1). The downstream region of the *tacA* ORF was PCR-amplified from *C. crescentus* gDNA with primers 7717/7718 (fragment 2). SOE-PCR was performed with the two fragments as template and primers 7925/7718. The resulting product was then digested with *Hind*III/*Eco*RI and ligated into pNPTS138 digested with the same enzymes.

pNPTS138-*tacA(D54E)*: The mutant allele was generated by SOE-PCR using *C. crescentus* gDNA as template and flanking primers 7958/7961 and mutagenic primers 7959/7960. The product was digested with *Pst*I/*Eco*RI and cloned into pNPTS138 digested with the same enzymes.

pNPTS138-Δ*spmX*: Two DNA fragments flanking the *spmX* ORF were PCR-amplified from *C. crescentus* gDNA. Fragment 1 was amplified with primers 7268/7269, fragment 2 was amplified with primers 7270/7271. The two fragments were joined by SOE-PCR using primers 7268/7271, digested with *Pst*I/*Eco*RI and ligated into pNPTS138 digested with the same enzymes.

pNPTS138-Δ*CC0168*: The region upstream of CC0168 was amplified from *C. crescentus* NA1000 gDNA with primer pairs 3490 and 3491. The PCR product was cloned into pGEM-T Easy, sequenced and cut out with *Hind*III and *Kpn*I. The region downstream of CC0168 was amplified from *C. crescentus* NA1000 gDNA with primer pairs 3492 and 3493. The PCR product was cloned into pGEM-T Easy, sequenced and cut out with *Bam*HI and *Kpn*I. Both cut PCR products were ligated in a triple ligation with pNPTS138 digested with *HindI*II and *Bam*HI.

pET28a-*His-shkA*: *shkA* was PCR-amplified from *C. crescentus* gDNA with primers 5988/5486, the product was digested with *Nde*I/*Hind*III and cloned into pET28a digested with the same enzymes.

pET28a-*His-shpA*: *shpA* was PCR-amplified from *C. crescentus* gDNA with primers 6015/6016, the product was digested with *Hind*III/*Eco*RI and cloned into pET28a digested with the same enzymes.

pET28a-*His-tacA-RD*: The sequence of *tacA* encoding the receiver domain was PCR-amplified from *C. crescentus* gDNA with primers 7925/6013, the product was digested with *Hind*III/*Eco*RI and cloned into pET28a digested with the same enzymes.

pET32b-*Trx-His-shkA-HK*: The sequence of *shkA* encoding the kinase catalytic core including the REC1 domain was PCR-amplified from *C. crescentus* gDNA with primers 6110/6011, the product was digested with *Hind*III/*Eco*RI and cloned into pET32b digested with the same enzymes.

pET32b-*Trx-His-shkA-RD2*: The sequence of *shkA* encoding the REC2 domain was PCR-amplified from *C. crescentus* gDNA with primers 6109/6009, the product was digested with *Hind*III/*Eco*RI and cloned into pET32b digested with the same enzymes.

pMT687-*tacA*: *tacA* was PCR-amplified from *C. crescentus* gDNA with primers 5487/5490, the product was digested with *Nde*I/*Kpn*I and cloned into pMT687 digested with the same enzymes.

pMT687-*tacA(D54E)*: The mutant allele was generated by SOE-PCR using *C. crescentus* gDNA as template and flanking primers 5487/5490 and mutagenic primers 5488/5489. The product was digested with *Nde*I/*Kpn*I and cloned into pNPTS138 digested with the same enzymes.

pQF-*PA5295*: *PA5295* was PCR-amplified from pRV-*PA5295* with primers 8974/8975, the product was digested with *Hind*III/*Kpn*I and cloned into pMT687 digested with the same enzymes.

pQF-*PA5295(AAL)*: *PA5295(AAL)* was PCR-amplified from pRV-*PA5295(AAL)* with primers 8974/8975, the product was digested with *Hind*III/*Kpn*I and cloned into pMT687 digested with the same enzymes.

pRKlac290-*spmX*: The promoter region of *spmX* was PCR-amplified from *C. crescentus* gDNA with primers 6864/6865. The resulting product was digested with *Eco*RI/*Pst*I and ligated into pRKlac290 digested with the same enzymes.

pRKlac290-*staR*: The promoter region of *staR* was PCR-amplified from *C. crescentus* gDNA with primers 6621/6622. The resulting product was digested with *Eco*RI/*Pst*I and ligated into pRKlac290 digested with the same enzymes.

pAH10: pUC19 was digested with *Hind*III and *Pci*I to integrate the following annealed oligos 5786/5903/5904/5905 downstream of the MCS.

pAH87: pMT375 was digested with *Xba*I and its insert was replaced with the *Nsi*I-*Pac*I cassette of pAH10 amplified with 7284 and 7287 digested with *Nhe*I/*Spe*I.

pAH99: *dendra2* was PCR-amplified from pDHL851 using primers 7503 and 7507, which also introduced a stop codon and the RBS of pQE70 upstream of *dendra2.* The product was digested with *Hind*III and *Pac*I and ligated into pAH87 digested with the same restriction enzymes.

pAH111: The promoter region of *spmX* was PCR-amplified from *C. crescentus* gDNA with primers 6864 and 7512, digested with *Eco*RI/*Hind*III and ligated into pAH99 digested with the same enzymes.

pAH139: *dendra2-ssrA* was generated by PCR from pDHL851 with primers 7503 and 6478. The resulting product was digested with *Hind*III/*Xho*I and ligated into pAH111 digested with the same enzymes.
